# Targeting an Olfactory Receptor Mitigates High Residual Platelet Reactivity and Arterial Thrombosis Through Actin Cytoskeleton Depolymerization

**DOI:** 10.1101/2024.09.30.615956

**Authors:** Anu Aggarwal, Vara Prasad VN Josyula, Nancy Wang, Moua Yang, Young Jun Shim, Quinn P. Kennedy, Reina Samuel, Naseer Sangwan, Suman Guntupalli, Matthew Godwin, Mariya Ali, Courtney Jennings, Bhairavi Rajasekar, Alliefair Scalise, Huijun Edelyn Park, Shaun Stauffer, Keith McCrea, Thomas McIntyre, Scott J. Cameron

## Abstract

**Background:** Despite antiplatelet therapy, some patients remain at high risk for ischemic events due to medication non-responsiveness or High Residual Platelet Reactivity (HRPR). Our goal was to target an orphan G-Protein-Coupled Receptor (GPCR) on platelets belonging to the olfactory receptor family as a new antithrombotic strategy.

**Methods:** Using an engineered reporter cell line expressing Olfactory Receptor 2L13 (OR2L13) that was recently identified as an orphan GPCR to limit platelet reactivity, we performed a high-throughput screen (HST) of non-odorant bioactive compounds with counter-screen validation, followed by studies to determine changes in platelet function in healthy subjects and in patients with coronary artery disease and peripheral artery disease. Phospho-proteomics identified key signal transduction pathways, and a variety of *ex vivo* assays and *in vivo* functional studies examined the impact of an identified non-odorant compound on platelet signaling, platelet biomechanics, and thrombosis.

**Results:** We identified 6 ligands specific for OR2L13 that function as receptor agonists, leading to the suppression of platelet aggregation and α-granule exocytosis via P2Y_12,_ PAR1, Thromboxane Receptor (TxR) and glycoprotein VI (GPVI) receptors, suggesting involvement of common downstream mediator. The lead OR2L13 agonist identified phosphorylates heat shock protein 27 (HSP27) and depolymerizes the platelet filamentous actin cytoskeleton which was further confirmed in a clot retraction assay (CCF0054500 clot area 70.6 vs. vehicle clot area 5.2, P<0.0001). This anti-thrombotic effect of the OR2L13 agonist was reversed by a HSP27 inhibitor (clot area 3.6, P<0.0001). In a murine cremaster arterial injury model, platelet accumulation at the injury site was reduced by 88.9% by the lead OR2L13 agonist compared to vehicle (P<0.0003) without altering fibrin generation *in vivo*, or interfering with the coagulation cascade, and without impairing the protective mechanism of platelet hemostasis.

**Conclusion:** We describe and characterize the first non-olfactory tool to target an olfactory receptor for the purpose of inhibiting platelet activation and thrombosis through downstream HSP27 in a comprehensive investigation using a first-of-its kind platelet inhibitor targeting an orphan platelet GPCR.

## Introduction

Cardiovascular disease remains a leading cause of mortality globally ^1^. Despite Goal-Directed Medical Therapy (GDMT), residual cardiovascular risk means that cardiovascular-related deaths continue to rise ^1–4^. Thrombosis is a feared and life-threatening consequence for patients with coronary artery disease (CAD) and peripheral artery disease (PAD). Clinical thrombosis progresses and ends with the activation of platelets by agonists through cell surface receptors (biochemical activation) and culminates in thrombus formation. One explanation for residual cardiovascular risk may be the unpredictable behavior of antiplatelet drugs and persistent platelet activation by shear stress (biomechanical activation) that current therapeutics cannot mitigate ^5,6^.

Biomechanical platelet activation alters the platelet phenotype and may lead to defects in the protective mechanism of hemostasis while permitting ongoing High Residual Platelet Reactivity (HRPR) which is a risk for thrombosis ^5,7^. In patients with established vascular disease, plaque formation in arteries leads to disturbed blood flow (D-Flow) which causes biomechanical platelet activation. We previously reported in humans and mice that platelet reactivity increases and persists after Myocardial Infarction (MI), highlighting an unmet need in the care of patients with thrombotic disease^8–10^.

The olfactory receptor (OR) family of GPCRs represents 18 gene families and 300 subfamilies that encode transmembrane G protein-coupled receptors (GPCR) on the cell surface. While ORs are primarily located in sensory neurons of the nasal epithelium, ORs are now known to be expressed ectopically in non-olfactory tissues such as airway smooth muscle, the kidney, colonic and vascular epithelium, and keratinocytes ^11–13^. Ectopic OR expression regulates important physiological and pathological processes, including blood pressure (BP), airway smooth muscle tone, vascular tone and remodeling^14–18^ that lead to ORs as emerging therapeutic targets for disease. Of the hundreds of ORs identified in humans, we identified only a few in platelet precursor megakaryocytes, and just 4 in healthy adult platelets ^19^. This presents an opportunity for targeted therapy for thrombotic diseases. Among the 4 ORs identified in adult platelets, OR2L13 expression is highest and stored in alpha granules prior to membrane surface translocation in platelets subjected to disturbed blood flow (D-flow) conditions^19^. A major objective of this study was evaluate OR2L13 as a druggable target using a high-throughput screen (HTS) and investigate the hitherto undiscovered signal transduction pathways in platelets to limit thrombosis through mechanisms unreachable by current antiplatelet drugs.

## Methods

### Human Subjects

Healthy subjects for different experiments were recruited only after informed consent by a coordinator not involved in clinical care using the following protocols: 19-1451 for patients and 20-413 for healthy volunteers. CAD and PAD patients were recruited from the clinic for the validation of the results. All study procedures were conducted in accordance with the Declaration of Helsinki and our approved local Institutional Review Board protocol.

### Experimental Animals

Murine protocols were approved by the University Committee on Animal Resources (UCAR). 6-8-week-old male and female wild-type C57BL/6J mice were used in this study ((Jackson laboratories, Bar Harbor, ME). Retro-orbital blood collected into heparinized Tyrodes solution, as described by us previously, was used to isolate mouse blood for complete blood counts and to keep platelets in a silent state prior to agonist stimulation ^10^. Complete blood counts mice were determined from retro-orbital blood into EDTA tubes on an Abaxis HM5C & VS2 automated complete blood analyzer (Allied Analytic, LLC, Tampa, FL).

### High Throughput Screen (HTS)

An OR2L13 HEK293 cAMP-Response-Element (CRE) reporter cell line was used to screen the 8K bioactive library (Life Chemicals, Inc. Niagara-on-the-Lake, Ontario, Canada). Using the robotic system (ECHO 550), 150nL of 10mM stock ligands were added to the 384 well plates to the final working concentration of 50μM. Then, 30K of OR2L13 HEK293 cAMP reporter cells were plated in 30μL phenol free DMEM media. In each sample plate, DMSO (vehicle), 500μM (-) Carvone (OR2L13 positive control) and 3 µM Forskolin (cAMP production positive control) wells were included. Plates were then incubated for 4 hours, after which 30μL of Steadylite Luciferase reagent (Perkin Elmer) was added to assess luciferase activity with a BioTek Cytation5 plate reader. The hits above 25% threshold of Forskolin (positive control) were further validated in HEK293 cAMP cells along with OR2L13 HEK293 cAMP in triplicates to confirm the specificity of ligands for receptor OR2L13. To calculate the half maximal effective concentration (EC_50_) for the top hits, dose response (195 nM to 100µM) was performed. The counter-screen validation experiment used the following concentrations for the 6 lead compounds: 1×10^−4^ M, 5×10^−5^ M, 2.5×10^−5^ M,1.25×10^−5^ M, 6.25×10^−6^ M, 3.13×10^−6^ M, 1.568×10^−6^ M, 7.88×10^−7^ M, 3.91×10^−7^ M, 1.9510^−7^ M.

### Validation of HTS hits using Dynamic Mass Redistribution on OR2L13-HA cAMP/PKA: Cells were

grown in DMEM media supplemented with Fetal Bovine Serum (FBS; 10%), and Pen-Strep (1%) in a humidified atmosphere at 37°C in 5% CO2. Prior to each Dynamic Mass Redistribution (DMR) experiment, 50,000 cells/well were seeded in a 96-well fibronectin-coated Corning Epic microplate in DMEM media (100 µL) and incubated overnight. The following morning, cells were washed with 1X HBSS buffer with HEPES (20 mM; pH 7.4) multiple times and the cells were resuspended in 1X HBSS buffer with HEPES. Next, the DMR plate was briefly centrifuged (100g for 15 second at room temperature) to allow the cells to settle at the bottom of the plate and then kept at room temperature for 1 hour. During measurements, basal DMR response was collected for 15 minutes to obtain a baseline which was defined as the zero point. For the DMR dose response experiments, different concentrations of the test compound of interest concentration) or vehicle were used and the DMR signal was read for 60-90 min in a Corning Epic BT system (Corning Epic).

### Off-target evaluation of lead compounds on other established drug targets

Reporter cell lines expressing GPCRs and ion channels were utilized to determine off-target effects of OR2L13 ligands (Eurofins DiscoverX, Sandiego CA). In brief, the assay read out determines whether putative OR2L13 ligands were agonists or antagonists for more than 70 targets using cAMP production, Ca^2+^ mobilization, ion flux, and nuclear receptor activity readouts. The readout provides potency (IC_50_, EC_50_) and efficacy (E_max_) from a multipoint concentration response-curve (CRC), the top concentration starting at 10 μM.

### Light Transmission Aggregometry

To determine the effect of the ligands on human platelets, Light Transmission Aggregometry (LTA) was used. Blood from healthy controls (n=7) was collected in sodium-citrate tubes, as described by us previously^19^. Within 30-60 minutes of collection, anticoagulated whole blood was centrifuged at 200g for 15 minutes at room temperature (RT) to obtain platelet rich plasma (PRP). Platelet Deplete plasma (PDP) was obtained from the remaining sample by re-centrifugation at 2500 g for 10 minutes and was used to initiate baseline optical density. PRP from healthy subjects was pre-treated with ligands used in a concentration of 100 µM or the purposes of the screen for 30 minutes at 37°C. Platelet aggregation was tested following stimulation with platelet agonist, Adenosine diphosphate (ADP) (1-5μM) where formation of platelet aggregates lead increased light transmission. For data analysis, the % aggregation in response to ADP was compared between ligands and vehicle (DMSO).

### Platelet Flow Cytometry

Ligands from the initial HTS were assessed for their platelet inhibitory effect by assaying surface P-Selectin expression through additional platelet receptors as a surrogate for α-granule exocytosis. Platelets were isolated from whole blood collected from healthy controls in sodium-citrate tubes. Whole blood was centrifuged at 200g for 15 minutes at RT and PRP was collected. Collected PRP was centrifuged again at 200g for 10 minutes to remove remaining WBCs and RBCs. Platelets were washed using Tyrode’s buffer at 1:1 ratio with the addition of Prostaglandin I2 (PGI_2_; 10nM). Then these washed platelets were resuspended in 1mL of Tyrode’s buffer. The washed platelets were pre-treated with ligands and vehicle (DMSO) for 30 mins. Then, 4mL of Tyrode’s buffer was added to make up the volume. After that, washed platelets were stimulated with the PAR 1agonist TRAP-6 (10μM), a thromboxane receptor agonist U46619 (5μM), a P2Y_12_ receptor agonist ADP (10μM) and a GPVI receptor agonist CRP (0.5μg/mL) for 15 minutes. Platelets were then incubated with phycoerythrin (PE)-conjugated anti-CD62P for 30 minutes at RT prior to fixation with 4% paraformaldehyde. Samples were analyzed using flow cytometry (BD Accuri C6 Plus) to acquire 10000 events. Data was analyzed using FlowJo software.

### Thrombosis *Ex-vivo* (biomechanical platelet activation)

Platelets are activated biomechanically by D-flow in atherosclerotic and aneurysmal arteries. To mimic these pathological processes, we used a T-TAS01 microfluidics system to determine the effect of lead OR2L13 ligands on biomechanically-induced thrombosis. As whole blood travels through collagen-coated capillaries under high shear (1500 s^−1^) or low shear (600 s^-1^), platelets adhere to the collagen surface and release ADP and Thromboxane A_2_. This leads to the formation and growth of the platelet thrombus. We tested the effect of lead compounds in whole blood from healthy subjects using PL (shear stress 1500 s^-1^) and AR chips (shear stress 600 s^-1^).

### Laser-injury thrombosis model

Thrombus formation in response to laser injury was measured in real-time as previously described^20,21^. FVB/NJ male mice (8-week-old) were obtained from the Jackson Laboratory and were treated with a daily dose of 75% w/v DMSO vehicle control (n=5) or 5 mg/kg CCF0054500 (n=5) intraperitoneally for three consecutive days. After the third treatment, the mice were anesthetized with an intraperitoneal injection of 125 mg/kg ketamine and 12.5 mg/kg xylazine. A patch of skin was removed from the neck and the jugular vein was cannulated with polyethylene tubing. The mice were intubated with polyethylene tubing for use with a respirator. The cremaster muscle was exteriorized and pinned onto a custom intravital microscopy tray. Platelet and fibrin accumulation were measured by infusing Dylight 488-labeled anti-platelet antibody (CD42b; 0.1 mg/kg body weight; Emfret Analytics) or Dylight 647-labeled anti-fibrin (clone 59D8; 0.3 mg/kg) monoclonal antibody through a jugular vein catheter. The cremaster muscle arterioles were injured using a MicroPoint Laser system (Andor, Belfast, UK). Data was acquired before and after laser injury using the brightfield, 488/520 nm, and 640/670 nm channel. Images were captured for 250 seconds at 2 frames/second using a CCD camera (ORCA Flash 4.0, Hamamatsu Photonics, Japan). Data was analyzed using Slidebook 6.0 (Intelligent Imaging Innovations, CO). Data from 29-31 thrombi were used to determine the median value of the integrated fluorescence intensity to account for variability in thrombus formation at any given experimental conditions. The integrated fluorescence intensity (RFU) for platelets and fibrin accumulation were calculated for each frame per the following equation: Integrated Fluorescence Intensity = Sum Intensity of the signal – (Average of the maximal background intensity) x Area in the pixel of the signal. The area under the curve (AUC) was calculated for individual thrombi using the trapezoid method with Igor Pro 9 (WaveMetrics, Inc., OR) and normalized to injury lengths to evaluate statistical significance. Injury lengths (µm) were determined in the brightfield channel as the distorted region along the luminal face of the vessel following the injury as previously described.

### IVC constriction thrombosis model

FVB/NJ male mice were obtained from the Jackson Laboratory and housed on a standard 12 h dark/light cycle. The mice were treated with a daily dose of 75% w/v DMSO vehicle control 5 mg/kg CCF0054500 intraperitoneally for three consecutive days. Following the third treatment, mice were anesthetized by continuous isoflurane induction (2%), with 100% oxygen at 2 L/min rate and placed on a heated pad at 37°C. The abdomen was incised, and intestines exteriorized to allow visualization of the inferior vena cava (IVC). All visible side branches of the IVC were ligated with 7-0 PROLENE sutures (Ethicon, Norderstedt, Germany) while back branches from the renal veins to the iliac bifurcation were cauterized. The IVC was then separated from the aorta and ligated in the same manner as the IVC side branches. The abdominal cavity was closed using 5-0 vicryl sutures (Ethicon) and adhesive skin glue. Mice were sacrificed 48 hours later and IVC thrombi were harvested, weighed, fixed by incubation in 4% (v/v) paraformaldehyde overnight at 4°C, and embedded in paraffin.

### Tail bleeding Assay

FVB/NJ male mice were obtained from Jackson Laboratories and housed on a standard 12 h dark/light cycle. The mice were treated with a daily dose of 75% w/v DMSO vehicle control or 5 mg/kg CCF0054500 intraperitoneally for three consecutive days. Following the third treatment, the mice were anesthetized with an intraperitoneal injection of 90 mg/kg ketamine and 10 mg/kg xylazine. A distal 10-mm segment of the tail was amputated with a scalpel followed by tail immersed in a pre-warmed isotonic saline solution in the 50-mL Falcon tube. Each animal was monitored for 9 minutes even if bleeding stopped, to detect any re-bleeding. Time to cessation of bleeding was evaluated in minutes followed by animal sacrifice with Ketamine and xylazine.

### Phosphokinase analysis

Unbiased phospho-proteomics deciphered the signaling pathway of OR2L13 ligands following lead compound incubation with platelets. Washed platelets from healthy subjects were pre-treated with DMSO or CCF0054500 (100µM) for 30 minutes at 37°C. Treated samples were processed and assessed by unbiased phosphoprotein analysis using liquid chromatography/mass spectrometry [LC-MS]). Briefly, 1.35mg protein from DMSO and CCF0054500 treated samples was digested with trypsin and then serine and threonine phosphorylation enrichment were performed using TiO_2_ based method. The overall abundance of peptides was moderate with over 16000 peptides identified, 33% of which were phosphorylated. The phospho-peptides identified in these samples were compared and three filters were used to identify differentially expressed phosphor-peptides with a p-value > 0.05. The results were also validated using Proteome Profiler Human Phospho-Kinase Array Kit (R&D Systems). For Phospho-Kinase Array, the washed platelets from healthy controls were treated with100µM CCF0054500 or vehicle (DMSO) for 30 minutes at 37°C. After that cells were lysed with lysis buffer and the assay was performed according to manufacturer’s recommendations. The results were confirmed with western blots for different proteins.

### Clot retraction *Ex vivo*

Washed platelets (2×10^8^ cells/mL) from healthy controls were incubated with CCF0054500 or vehicle (DMSO) for 30 minutes at 37°C with and without preincubation with 60µM BVDU (Brivudine) for 45 minutes at 37°C. Fibrinogen was added at a final concentration of 1 mg/mL with calcium chloride at final concentration of 2 mM. Thrombin (1U/mL) was added and then the tubes were incubated at 37°C. Images were captured in every 10 minutes until 1 hour and data presented as clot area.

### Statistics

Data are presented as the mean ± SEM or mean mean ± SD depending on distribution and population size unless e stated. Normality were evaluated by the Shapiro-Wilk test. For normally distributed data between 2 comparative groups, a 2-tailed Student’s *t* test was used. For nonparametric data, the Mann-Whitney *U* test was used. For Gaussian-distributed data in 3 or more groups, 1-way ANOVA followed by Bonferroni’s multiple-comparison test was used, otherwise the Kruskal-Wallis test followed by Dunn’s post-test correction was used. Significance was accepted as a *P* value of less than 0.05. Analyses were conducted using GraphPad Prism 7 (GraphPad Software).

## RESULTS

### Non-olfactory compounds activate olfactory receptor OR2L13

We previously reported that OR2L13 activation in platelets increases cAMP through odorant stimulation by a molecule that is the active ingredient in spearmint ^19^. Using an engineered reporter cell line activated by OR agonists, cyclic AMP (cAMP) production is readily detectable upon OR2L13 stimulation. We screened a library of non-odorant bioactive compounds known to activate GPCRs to identify and test a new approach for antiplatelet therapy. The 8K bioactive library screen identified 169 preliminary hits (2.1% hit rate) exceeding the 25% threshold Z’ which is the statistical separation between positive and negative controls in an assay than was greater than 0.5 for each plate in the HTS. To limit variability with cell passaging, data were normalized to forskolin in each plate (positive control, directly activates adenylyl cyclase to synthesize cAMP). In a counter-screen at 50 μM concentration with a vector HEK293-CRE cell line, we validated the specificity of those hits for OR2L13. 12 compounds emerged as agonists for OR2L13 (CCF0051970, CCF0054500, CCF0057537, CCF0054432, CCF0053625, CCF0052884, CCF0053070, CCF0058399, CCF0056873, CCF0052249, CCF0058334, and CCF0053066) (**Figure 1**).

**Figure 1:**
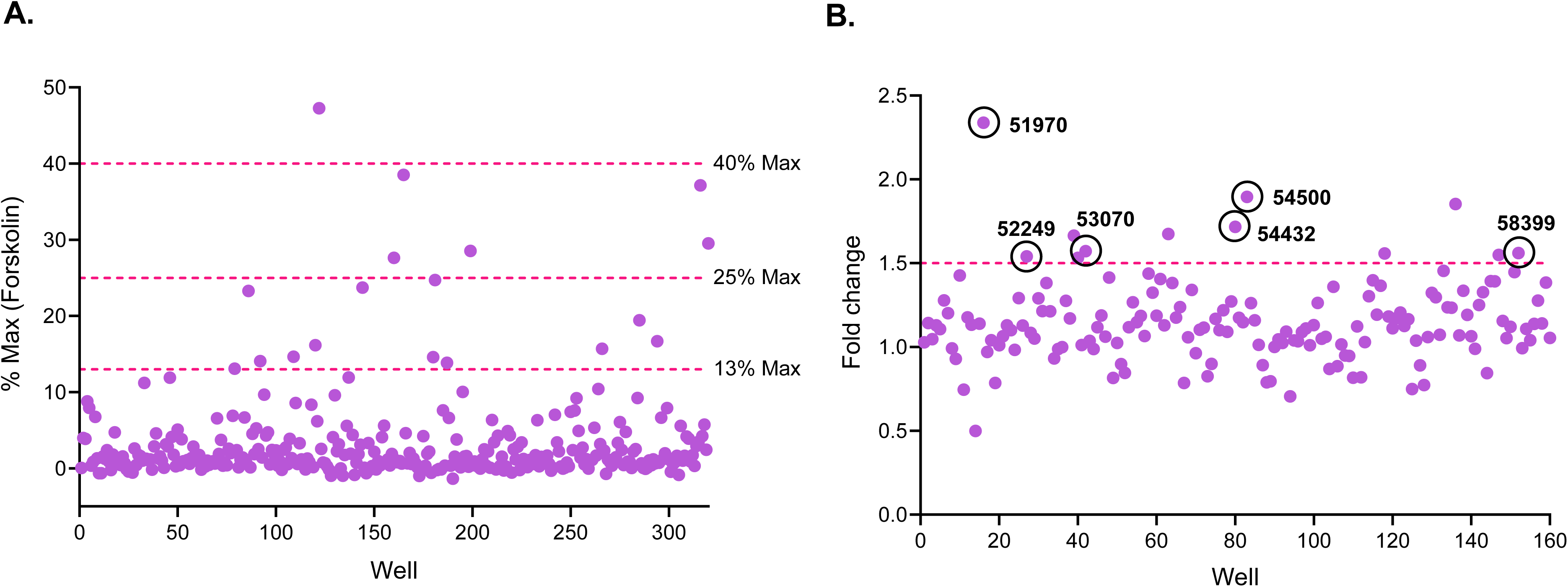
Screening of 8000 bioactive library for a non-odorant OR2L13 agonists. **(A**) Primary Screen. 8000 non-olfactory compounds using a OR2L13 HEK293 cAMP reporter cells. 169 hits (2.1% hit rate) >25% threshold were determined. All plates passed a Z’ >0.5. L-Carvone was used as a positive control for OR2L13 receptor and Forskolin as a positive control for adenylyl cyclase. **(B)** Counter-screen using a HEK293 cAMP reporter cell line was set up. Fold change of 169 hits was determined as the luminescence ratio of OR2L13 HEK293 cAMP/HEK293 cAMP reporter cell lines. Hits >1.5-fold above baseline (CCF0054500, CCF0054432, CCF0053070, CCF0052249, CCF0051970 and CCF0058399, each 50 µM final concentration) encircled in red were selected for further analysis.

Each of the 12 compounds demonstrated reporter luminescence >1.5-fold above baseline. The EC_50_ for each OR2L13 agonist was determined for only 6 ligands specific for OR2L13 (CCF0054500, CCF0054432, CCF0053070, CCF0052249, CCF0051970 and CCF0058399) as the other 6 compounds were unavailable in sufficient quantities (**Figure S1**). The chemical structures of the 6 lead compounds shared similar core and side chain features (**Figure 2**). Given that ORL13 is a GPCR, activation was further assessed by dynamic mass redistribution (DMR) for each lead OR2L13 compound in a dose-response manner. We found significantly higher GPCR responses at all doses with these compounds compared to vehicle (**Figure S2**). Collectively, these data validated 6 non-olfactory ligands as agonists for human OR2L13.

**Figure 2:**
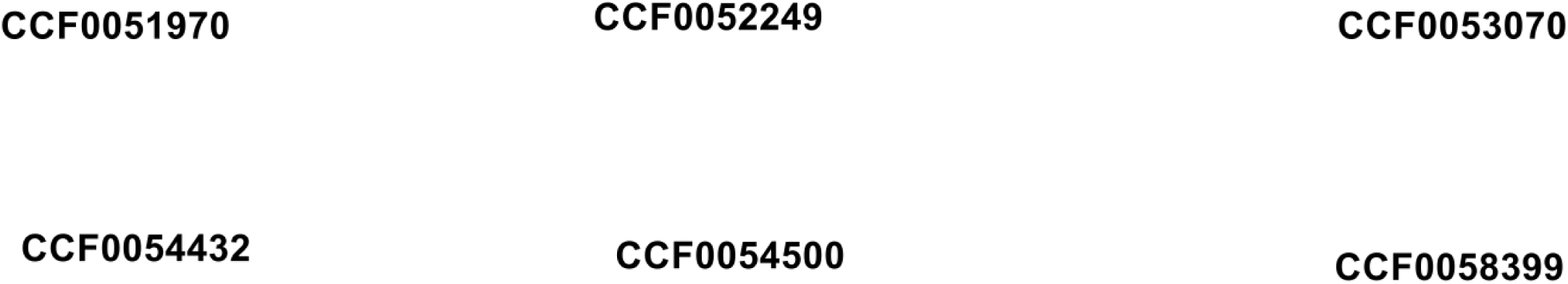
Lead OR2L13 agonists. The name of each led compounds is indicated.

### The effect of non-olfactory ligands of OR2L13 on platelet function

The efficacy of the 6 leading non-olfactory ligands to inhibit platelet function was determined in platelet rich plasma (PRP) and in individual, washed platelets. Light Transmission Aggregometry (LTA) determined platelet activation firstly by aggregation in PRP. The impact of 6 lead compounds as antiplatelet compounds was assessed following ADP-mediated platelet aggregation. Only three compunds (CCF0054500, CCF0052249 and CCF0058399) inhibited ADP-mediated platelet aggregation in healthy humans beyond 30% (**Figure 3A-B**). Antiplatelet properties of the three leading OR2L13 compounds were further evaluated by flow cytometry by the appearance of surface CD62-P (p-selectin) which signifies platelet activation by α-granule exocytosis. Evaluating platelet activation by α-granule exocytosis is an efficient manner that interrogates multiple platelet receptor-mediated activation pathways simultaneously (PAR1, P2Y_12_, thromboxane receptor, and glycoprotein VI). CCF0052249 and CCF0058399 did not suppress all platelet activation pathways in the manner observed with CCF0054500 (**Figure S3**). For this reason, CCF0054500 was progressed as a chemical probe for OR2L13, revealing an IC_50_ in the low μM range in to inhibit platelet aggregation in PRP from healthy subjects following platelet activation by ADP (**Figure 3C**). Surprisingly, CCF0054500 inhibited agonist-mediated platelet activation through all GPCRs assessed including those involving Gαq and Gαi, as well as through GPVI, suggesting a non-canonical downstream and common OR signaling mediator (**Figure 3D**). CCF0054500 was therefore pursued as our lead compound for mechanistic evaluation.

**Figure 3:**
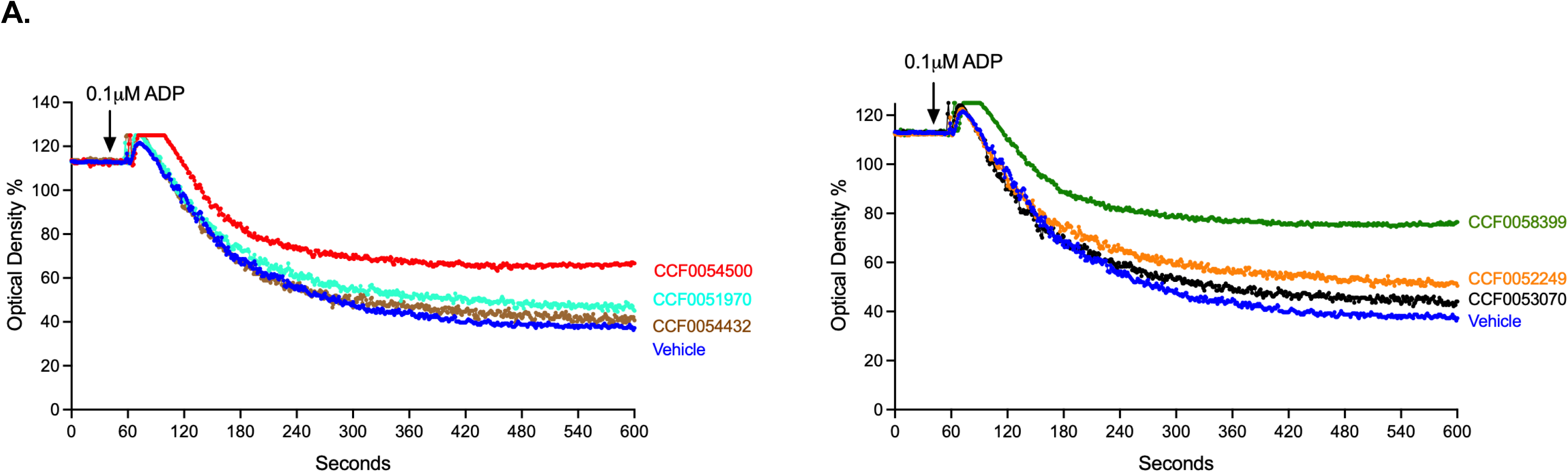
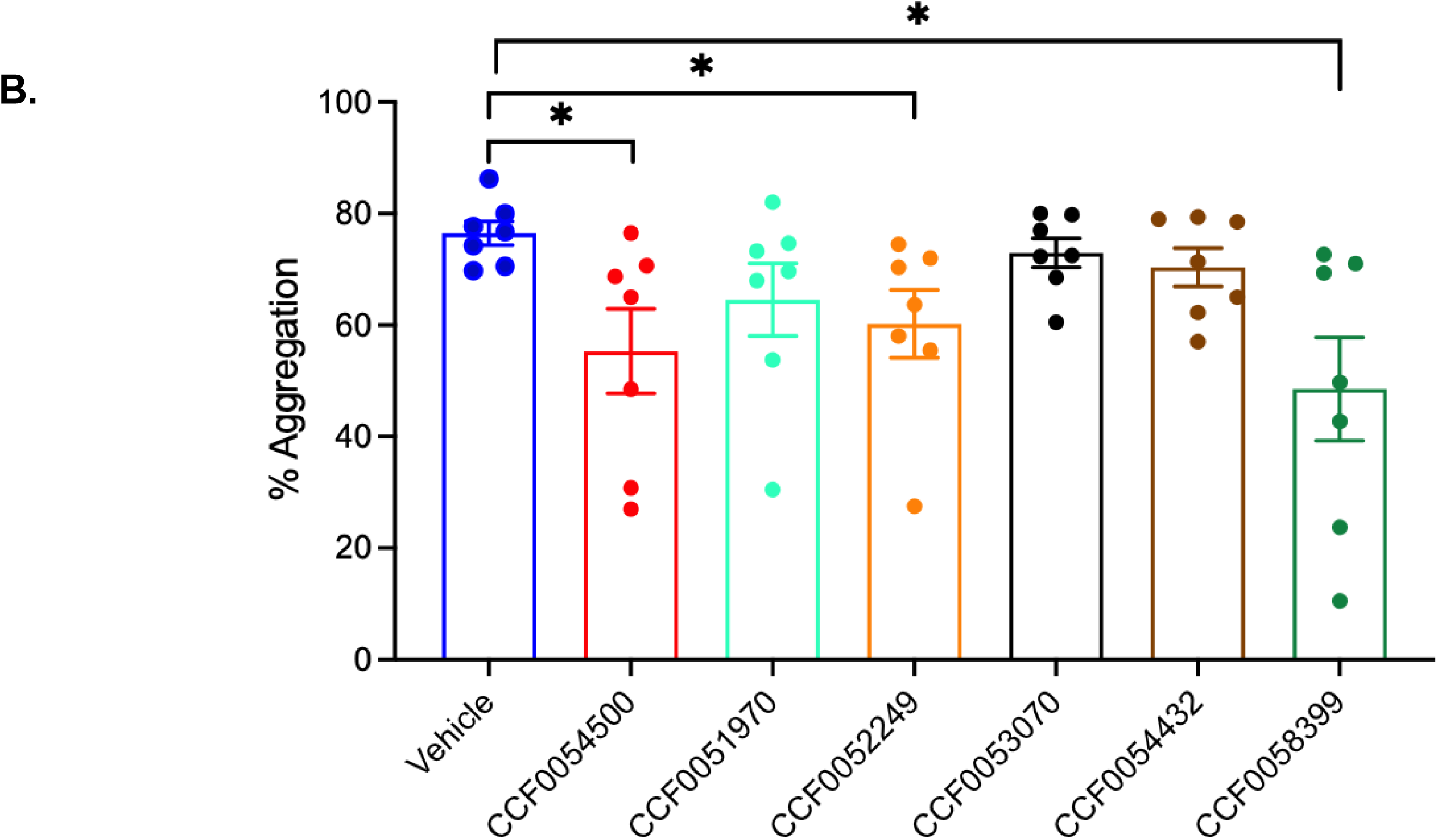
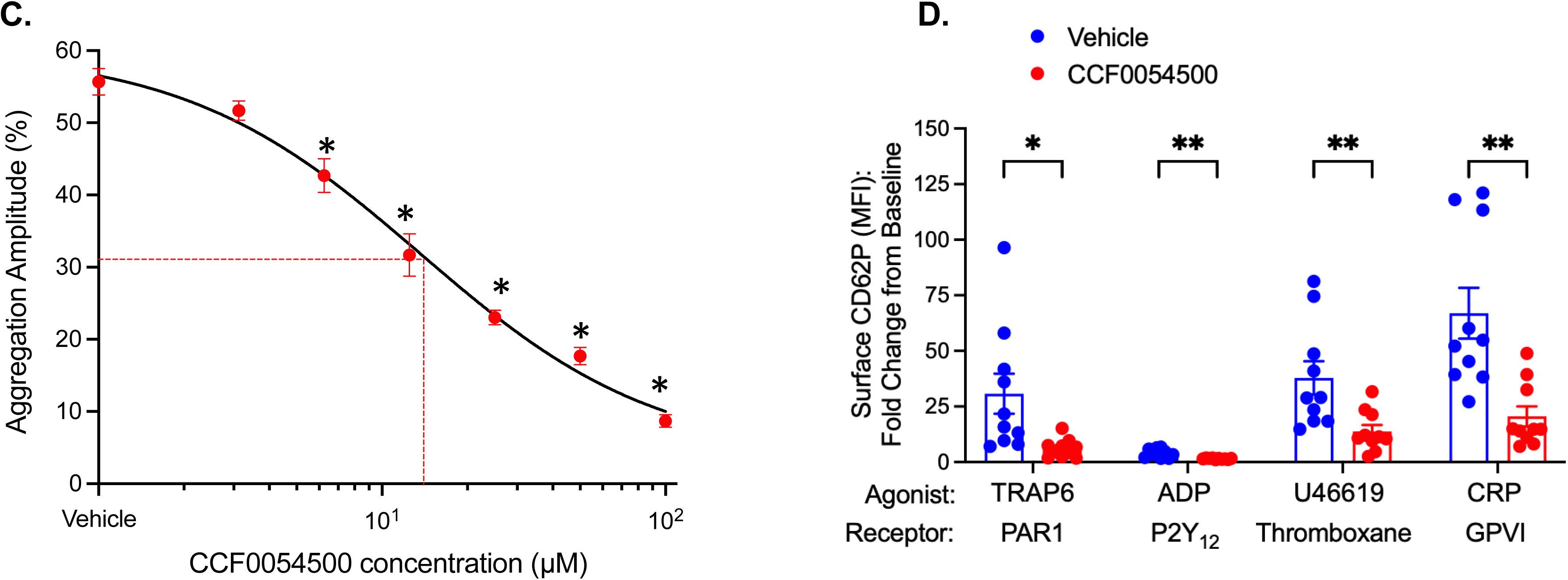
Impact of non-olfactory OR2L13 agonists on activation platelets from healthy humans. **(A)** Light Transmission Aggregometry for the 6 OR2L13 lead compounds (CCF0054500, CCF0051970, CCF0052249, CCF0053070, CCF0054432 and CCF0058399). Tracings are representative of Platelet Rich Plasma (PRP) 7 from healthy subjects. PRP was pre-incubated with 100μM of ligand or vehicle for 30 minutes at 37°C. After 30 seconds, 0.1μM ADP agonist was added to determine platelet aggregation. **(B)** Summarized aggregometry data are shown as mean % maximum aggregation ± SEM form n=7 independent healthy volunteers in each group, each performed in duplicate. *P* values <0.05 (∗).**(C)** Light Transmission Aggregometry dose response using the OR2L13 agonist antiplatelet compound CCF0054500 incubated in Platelet Rich Plasma (PRP) from healthy controls (n=7, each performed in duplicate). PRP was pre-incubated with increasing concentration of CCF0054500 compared to vehicle for 30 minutes at 37°C and followed by platelet activation with 0.1μM ADP. Summarized aggregometry data is shows as mean % aggregation ± SEM, n=3 independent healthy volunteers in each group, each performed in duplicate. The IC_50_ concentration of CCF0054500 for platelet inhibition is indicated by the red line. **(D)** Change in platelet surface P-selectin expression to signify platelet activation by alpha granule exocytosis. Washed platelets treated with agonists or vehicle for 30 minutes at 37°C. The results are expressed as Mean Fluorescence of P-selectin ± SEM (n=10, each performed in quadruplicate, t-test). TRAP6=Thrombin Receptor Activator for Peptide 6, ADP=Adenosine Diphosphate, CRP=collagen-related peptide. *P* values <0.05 (∗) and <0.01 (∗∗).

### OR2L13 agonist inhibits in-vivo thrombus formation under shear stress without affecting coagulation

CCF0054500 was reintroduced into whole blood from healthy subjects to assess thrombosis under shear stress conditions using the T-TAS01 system in which blood is passed through collagen-coated microfluidics channels. CCF0054500 decreased thrombosis in whole blood by >50% compared to vehicle, uniquely under conditions of high arterial shear rate (1500 s^−1^, **Figure 4A**) but not lower arterial shear rate (600 s^−1^, **Figure S4**). Given that CCF005400 inhibits thrombosis in whole blood, in isolated platelets, and in PRP from humans, assessing the impact on the coagulation cascade was prioritized in anticipation of finding bleeding as a side-effect. Platelet-deplete plasma (PDP) was used to assess real-time fibrin (Factor Ia) formation which is the most downstream component of the coagulation cascade. However, we did not observe any impact of CCF0054500 on fibrin formation in PDP isolated from healthy humans when assessed both by the rate of clot growth *ex vivo* or absolute clot size (**Figure S5**).

**Figure 4.**
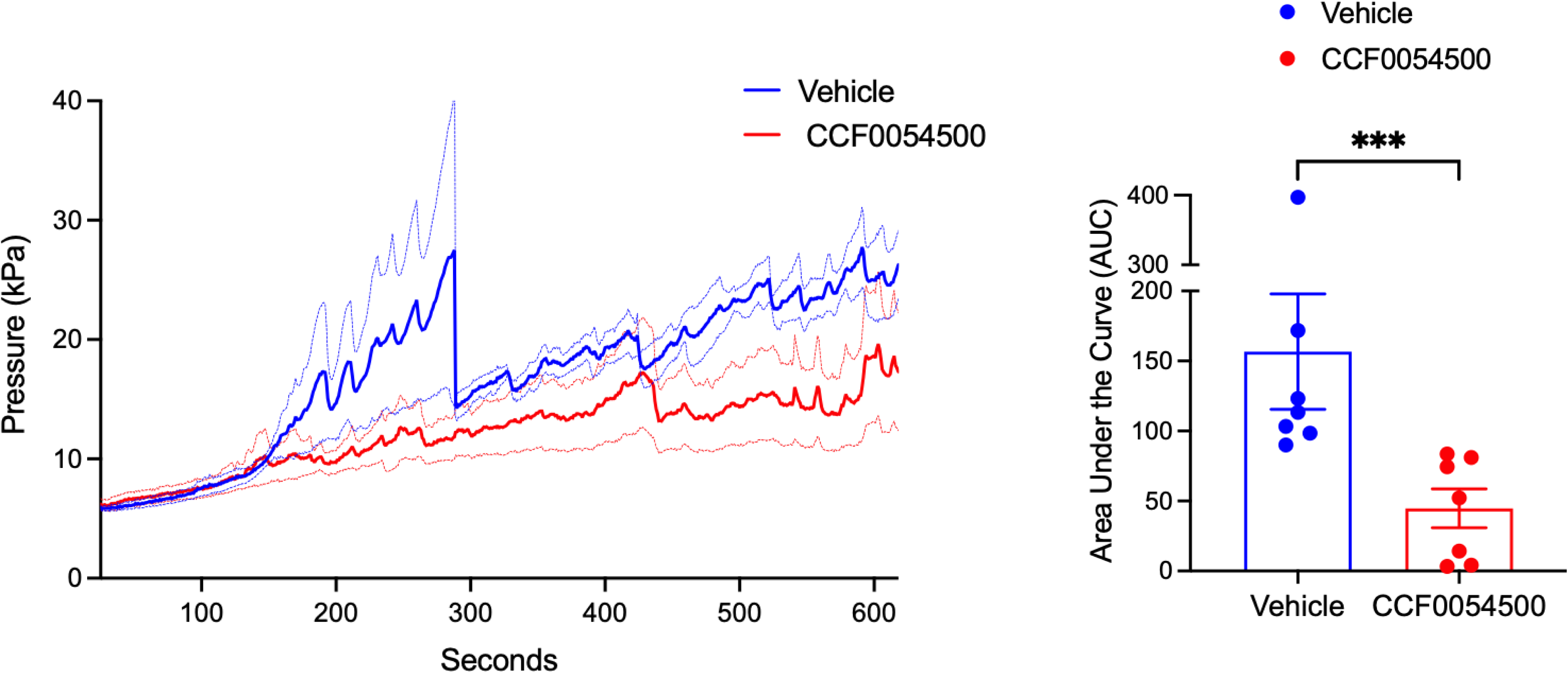
Impact of non-olfactory OR2L13 agonists lead compound on biomechanical platelet activation under high arterial shear stress. T-TAS® (Total Thrombus-formation Analysis System) evaluates thrombosis of whole blood under high arterial shear stress conditions in a microfluidics system (shear rate 1500.Sec^−1^) as a function of time as mean peak pressure ± SEM (broken lines). CCF0054500 decreases thrombotic occlusion of channel under shear stress compared with vehicle and summarized as Area Under the Curve (AUC, n=7 healthy volunteers). *P* values <0.001 (∗∗∗).

### OR2L13 agonist inhibits arterial thrombosis in-vivo

To ascertain the bioavailability of CCF0054500 and to assess cross-species efficacy, the compound was introduced into FVB/NJ mice by daily intraperitoneal (IP) injections to assess the impact on hemostasis and thrombosis *in vivo.* Laser injury-initiated thrombus formation in the mouse cremaster vasculature was measured by platelet and fibrin accumulation to the intima over time following consecutive daily doses of CCF0054500 (5 mg/kg i.p.). CCF0054500 decreased platelet accumulation by 88.9% compared to the vehicle control treatment group (**Figure 5A-C**). Similar to what was observed in humans, fibrin formation was unaffected by CCF0054500 (p = 0.54; **Figure 5D**) suggesting no impact on the coagulation cascade *in vivo* in mice. The median injury sizes were similar between the two treatments (**Figure 5E-F**), indicating that the differences observed were not from differences in the extent of vascular injury. These data suggest CCF0054500 prevents platelet accumulation in thrombus formation without an impact on fibrin formation which is consistent with what was observed in human platelet-deplete plasma *ex vivo* (**Figure S5**).

**Figure 5.**
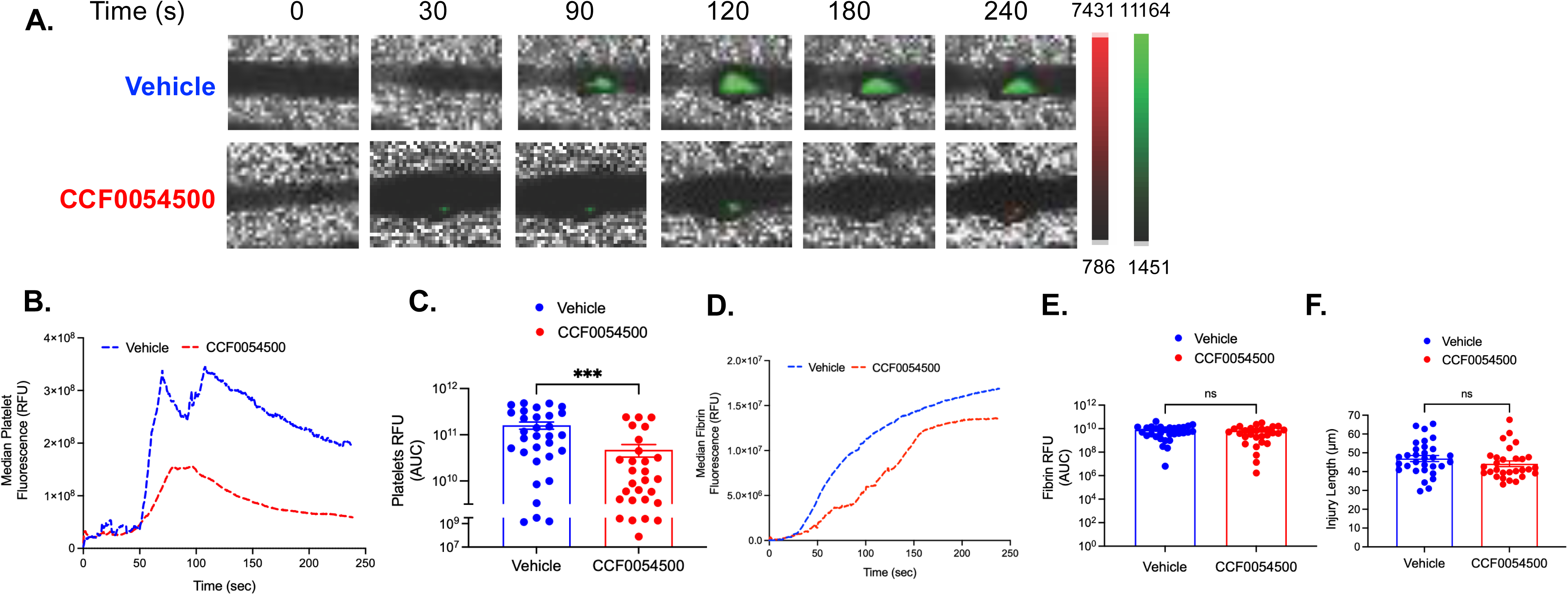
CCF0054500 decreases platelet accumulation *in vivo* in an arterial thrombus after laser injury. (A). Intravital microscopy images of the mouse cremaster arterioles showing platelet (green) and fibrin (red) accumulation following laser injury in vehicle and CCF0054500 (5 mg/kg x 3 days i.p.) treated groups. (B) Mean integrated fluorescence intensity over time of platelets. (C) Quantification of the normalized platelet accumulation with the median shown as mean ± SEM (D). Median integrated fluorescence intensity over time of fibrin (E). Quantification of the normalized fibrin accumulation as mean ± SEM. (F). Injury sizes between vehicle and CCF0054500 as mean ± SEM. Vehicle, N = 31 injuries from 4 mice; CCF0054500, N = 29 injuries from 3 mice. *P*-values were determined by non-parametric Mann-Whitney test. *P* values <0.001 (∗∗∗).

### OR2L13 agonist inhibits thrombosis in vivo under conditions of vascular obstruction

Mechanical stress on blood leads to thrombosis *in vivo* and is readily assessed by inferior vena cava (IVC) constriction. IVC constriction creates D-flow proximal to the suture line and exposes blood to external mechanical forces as well as stasis ^22^. FVB/NJ mice were treated with CCF0054500 (5 mg/kg daily i.p. for 3 days) followed by IVC constriction. CCF0054500-treated mice developed thrombi that were 49% smaller than vehicle-treated mice (weight 8.20±2.6 mg vs. 15.98±5.0 mg, p=0.0074) (**Figure 6A**). Thrombus composition of red blood cells (CD235a), platelets (CD41), white blood cells (CD45), and fibrin were similar in mice treated with CCF0054500 compared with vehicle (**Figure S6**). Baseline blood counts in mice were unaltered by CCF0054500 when given to mice illustrating the absence of toxic effects on hematopoiesis (**Figure S7**). Finally, CCF0054500 did not impact hemostasis given the duration of bleeding in CCF0054500-treated mice after tail amputation was similar to vehicle administration (**Figure 6B**).

**Figure 6.**
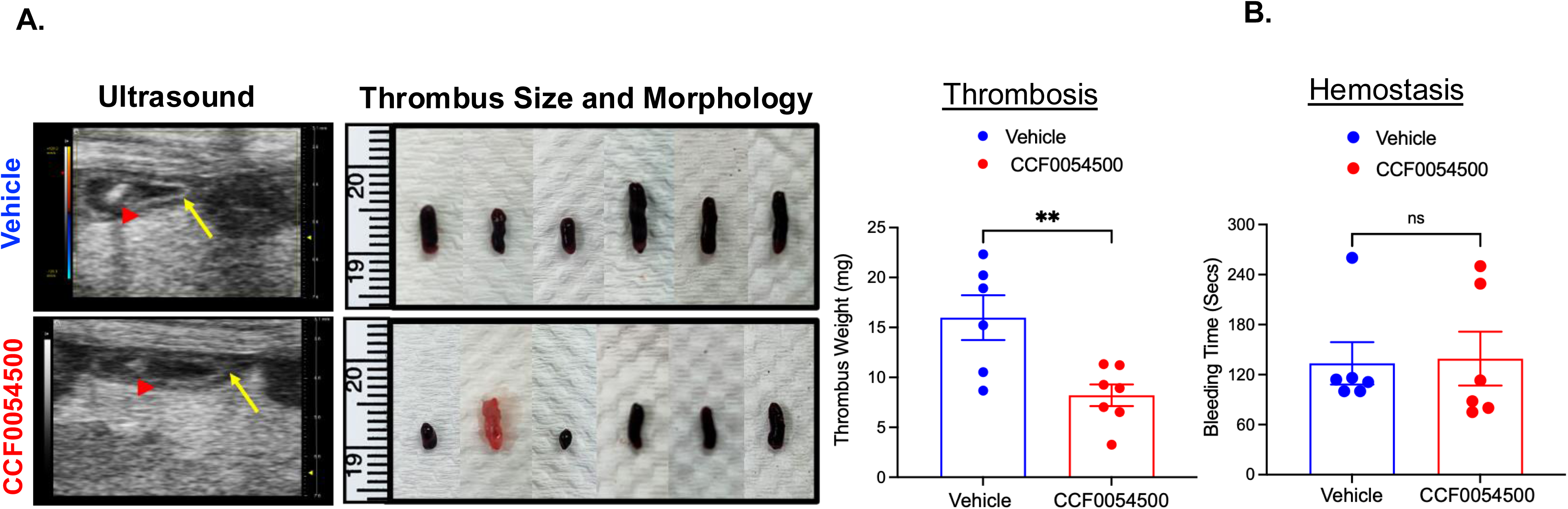
CCF0054500 decreases thrombus size *in vivo* in an inferior vena cava stasis model. **(A)** Smaller thrombus was detected by B-mode ultrasound (read arrowhead) in the group treated with CCF0054500 (5 mg/Kg/day x 3 days i.p.) than the vehicle-treated group by IVC constriction (yellow arrow is the region of ligature). Thrombus weight is represented as mean ± SEM (n=6-7, t-test). **(B)** CCF0054500 (5 mg/Kg/day x 3 days i.p.) does not alter hemostasis, indicated by tail bleeding times following tip amputation. Data are represented as mean time to cessation of blood flow ± SEM (n=6, t-test). *P* values <0.01 (∗∗).

### OR2L13 agonist suppresses high residual platelet reactivity in patients with vascular disease

Isolated, washed platelets from patients with coronary artery disease (CAD) or peripheral artery disease (PAD) were assessed for their ability to activate through four major platelet receptors with and without exogenous CCF0054500 incubation. Platelet reactivity through most surface receptors, especially in patients with PAD, and particularly through GPVI for both CAD and PAD, persisted despite antiplatelet therapy adherence (**Figure 7**). Irrespective of whether the patient was taking mono-antiplatelet therapy (aspirin or clopidogrel alone) or dual antiplatelet therapy (aspirin and clopidogrel together), the addition of CCF0054500 suppressed HRPR through all signaling pathways. This suggests a non-canonical pathway downstream of OR2L13 as a common point of convergence that disrupts platelet activation through the 4 surface receptor pathways assessed.

**Figure 7:**
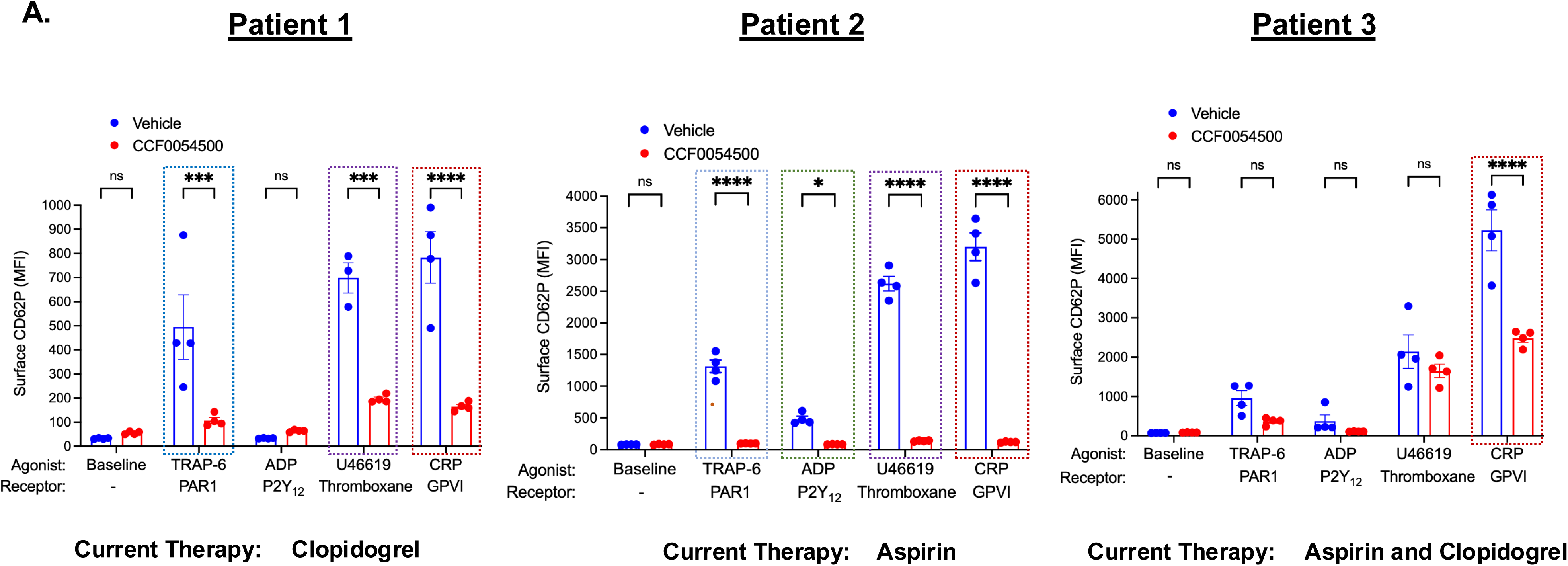
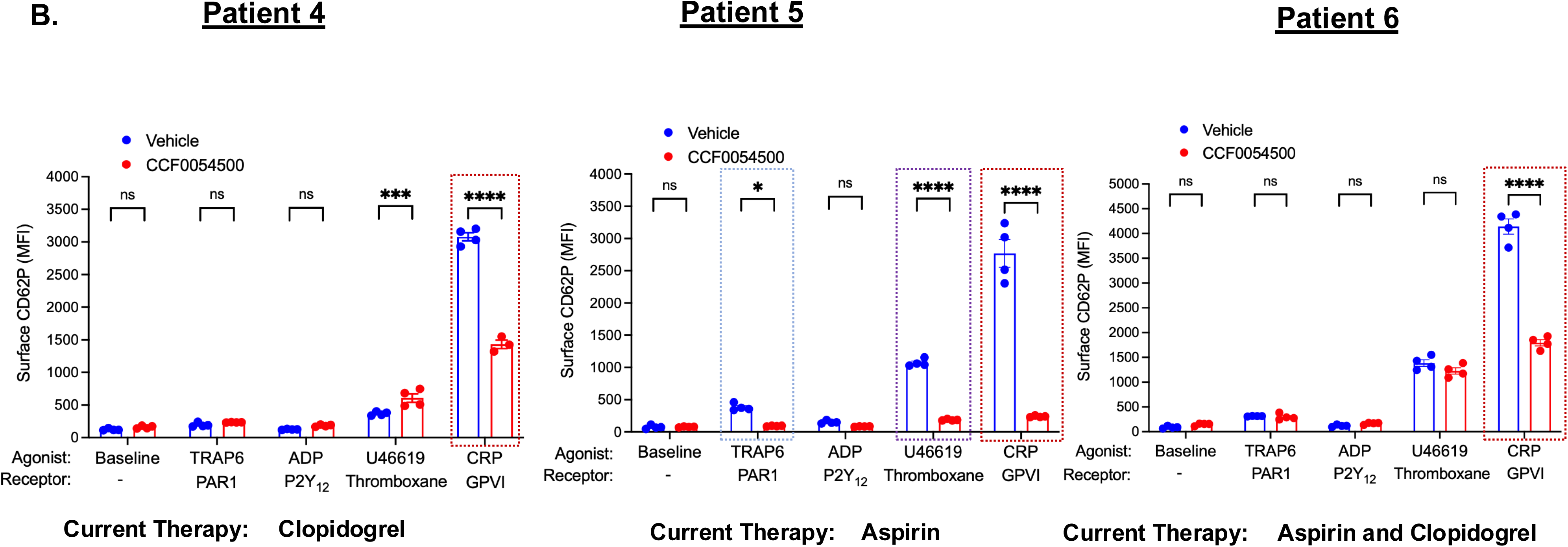
HRPR mitigated by CCF0054500 in patients with CAD and PAD on anti-platelet therapy. **(A)** Isolated platelets from patients with atherosclerotic PAD treated with CCF0054500 and Vehicle for 30 mins. Platelet reactivity in response to surface receptors agonists for each of 3 patients by technical quadruplicate. HRPR is prominent through the glycoprotein VI (GPVI) receptor in all patients (red box). HRPR is noted through the P2Y_12_ receptor (green box), thromboxane receptor (purple box), and PAR 1 (blue box). CCF0054500 suppress HRPR in all patients. *P* values < 0.05 (∗), 0.01 (∗∗), 0.001 (∗∗∗), and 0.0001 (∗∗∗∗). **(B)** Isolated platelets from patients with atherosclerotic CAD treated with CCF0054500 and Vehicle for 30 mins. Platelet reactivity in response to surface receptors agonists for each of 3 patients by technical quadruplicate. HRPR is prominent through glycoprotein VI (GPVI) receptor in all patients (red box). HRPR is noted through the P2Y_12_ receptor (green box), thromboxane receptor (purple box), and PAR 1 (blue box). CCF0054500 suppress HRPR in all patients. *P* values <0.05 (∗), 0.01 (∗∗), 0.001 (∗∗∗), and 0.0001 (∗∗∗∗).

### CCF0054500 activates platelet HSP27 and depolymerizes filamentous actin cytoskeleton

Platelet activation almost always triggers a cascade of protein kinases and causes protein phosphorylation events that alter platelet reactivity^23^. To identify a common downstream mediator from all platelet receptors that explains the antiplatelet effect of CCF0054500, we first utilized an unbiased approach to examine the platelet phosphoproteome by mass spectrometry. These studies revealed that CCF0054500 regulates proteins involved in cytoskeleton organization, actin filament depolymerization, and regulation of platelet organelles in human platelets (**Figure 8A**). We next used a targeted phosphokinase array which revealed the leading protein phosphorylated after platelet stimulation by CCF0054500 was heat shock protein 27 (HSP), culminating in phosphorylation of HSP27 residues Ser78, and Ser82 (**Figure 8B-C**). This observation was further validated in separate experiments in a different population of human subjects by performing immunoblotting with anti-HSP27 antibodies recognizing Ser15, S1er78 and Ser82 (**Figure 8D**).

**Figure 8:**
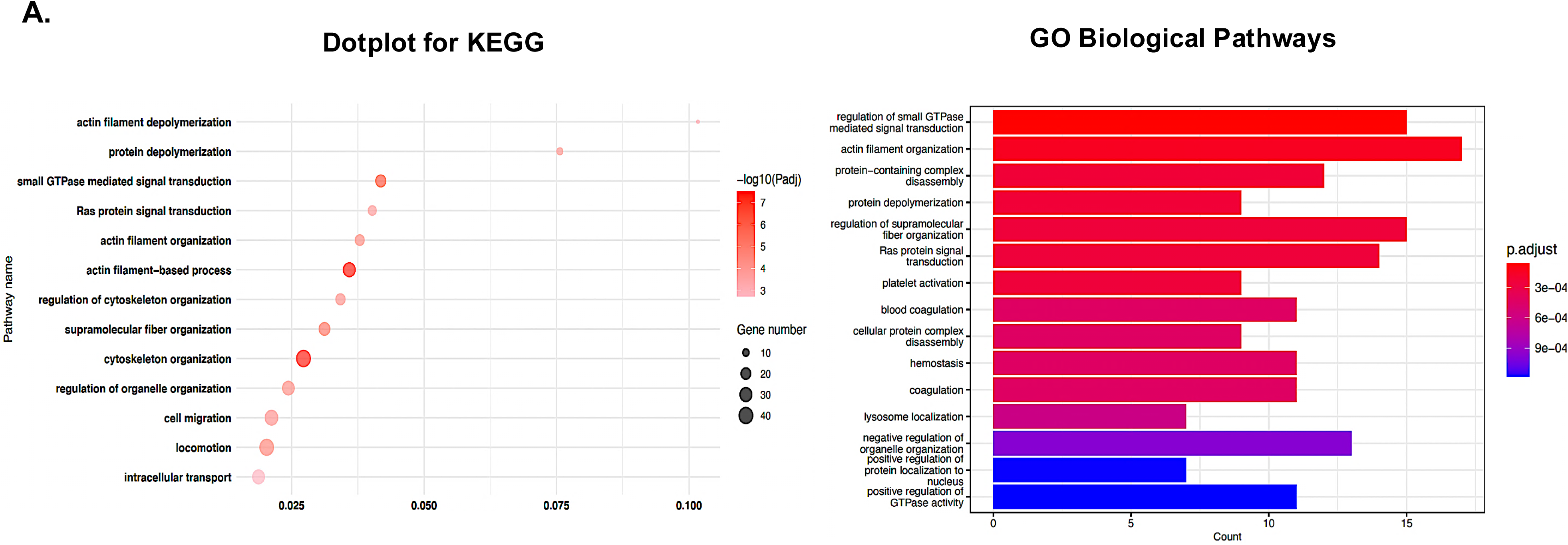
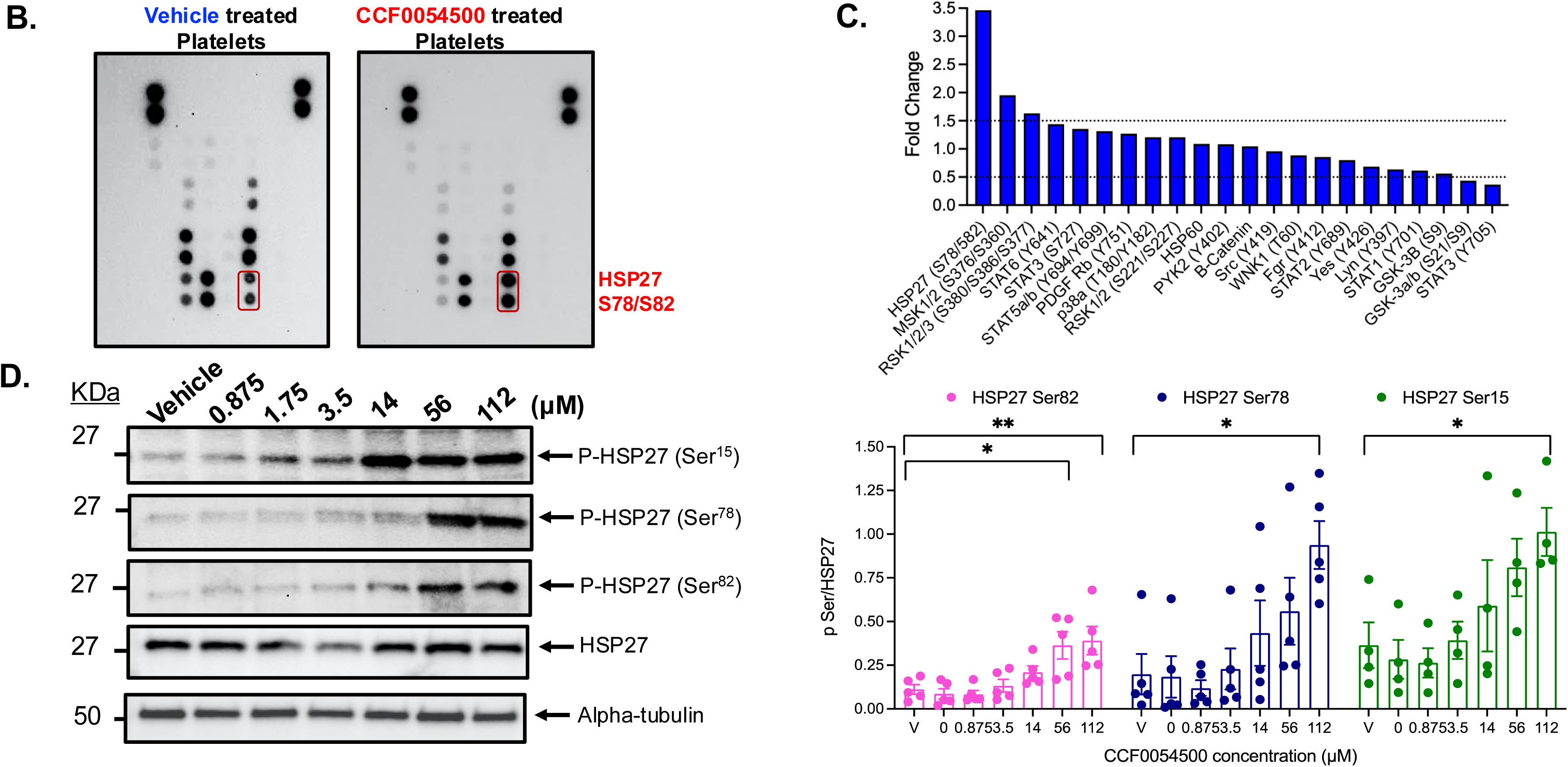
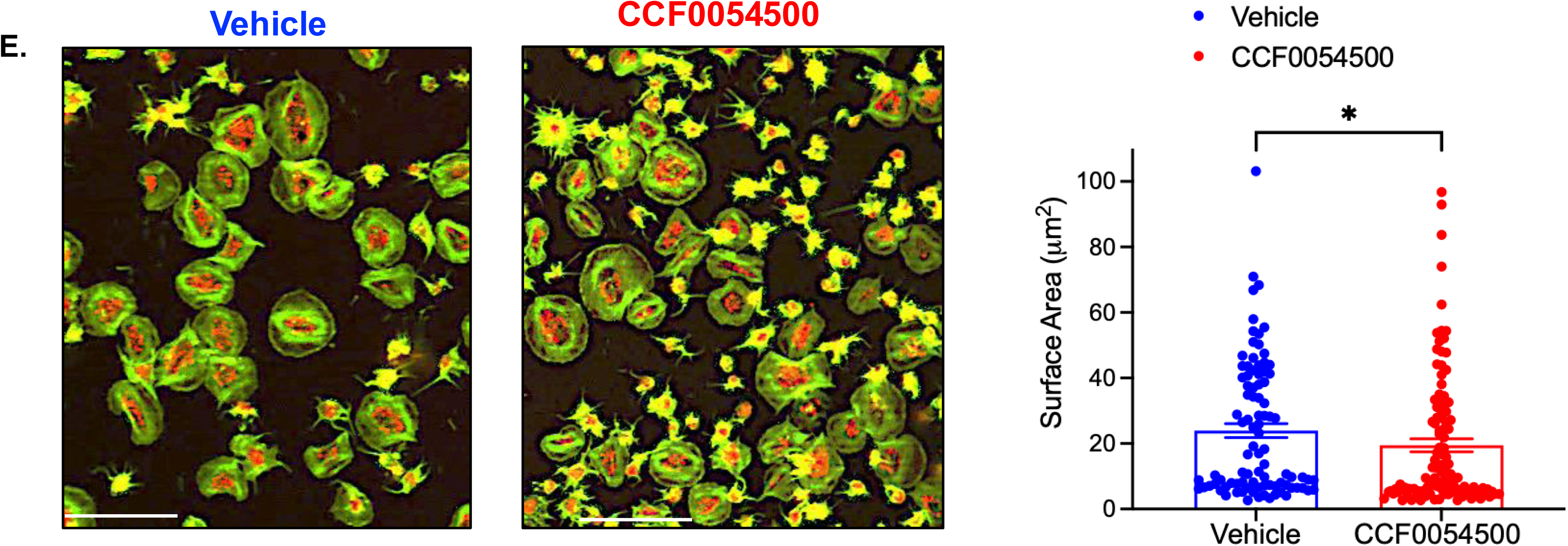
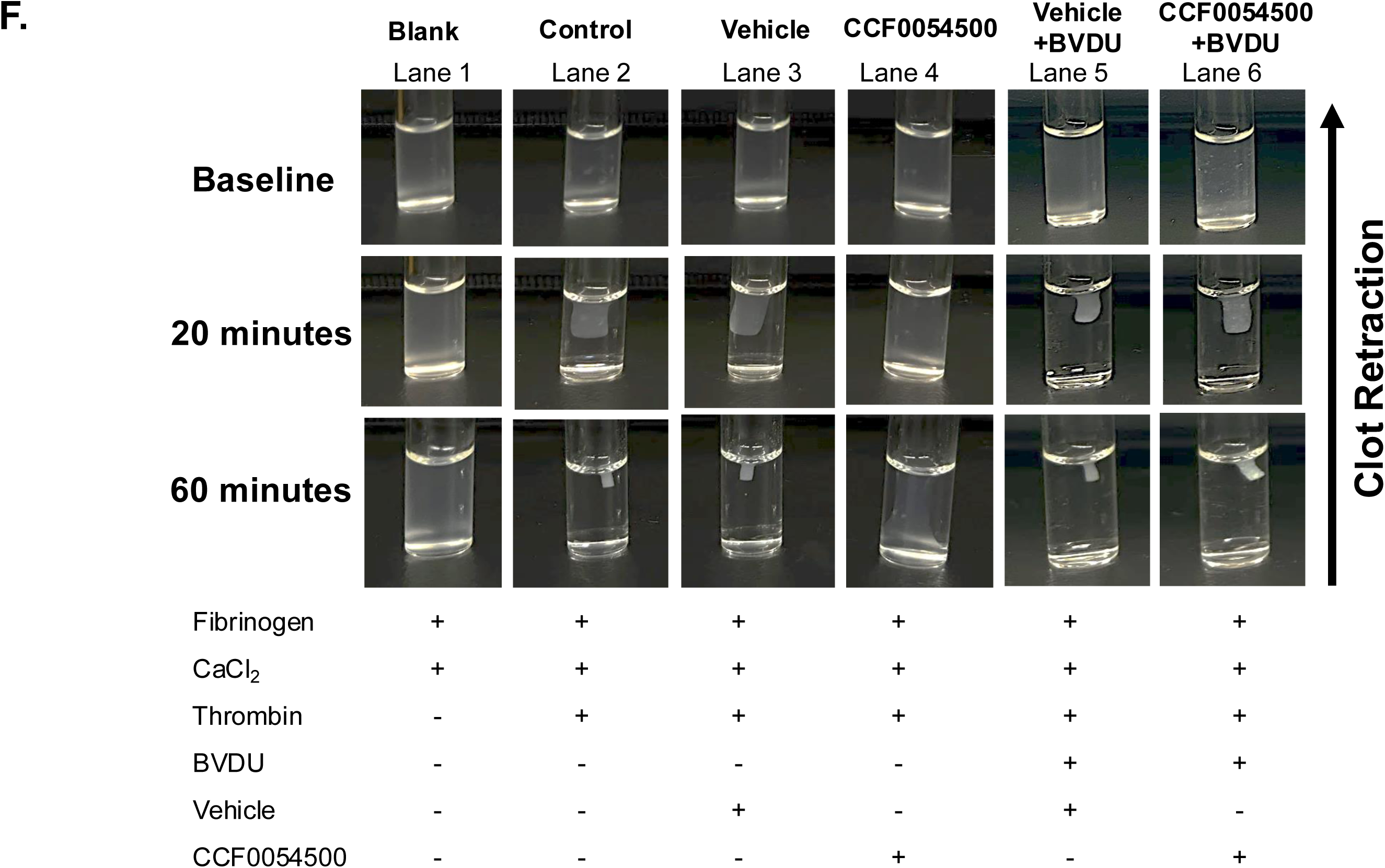
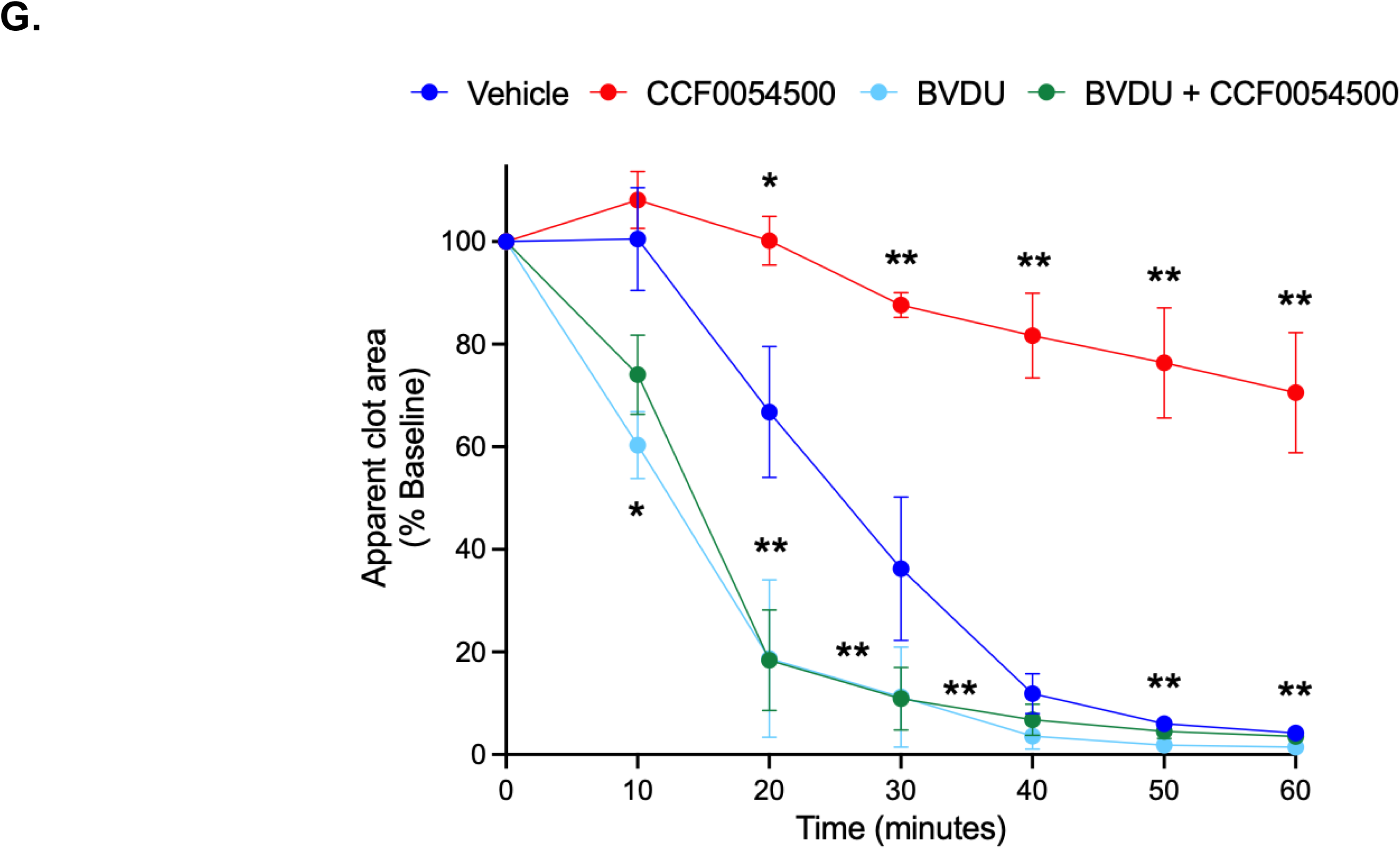
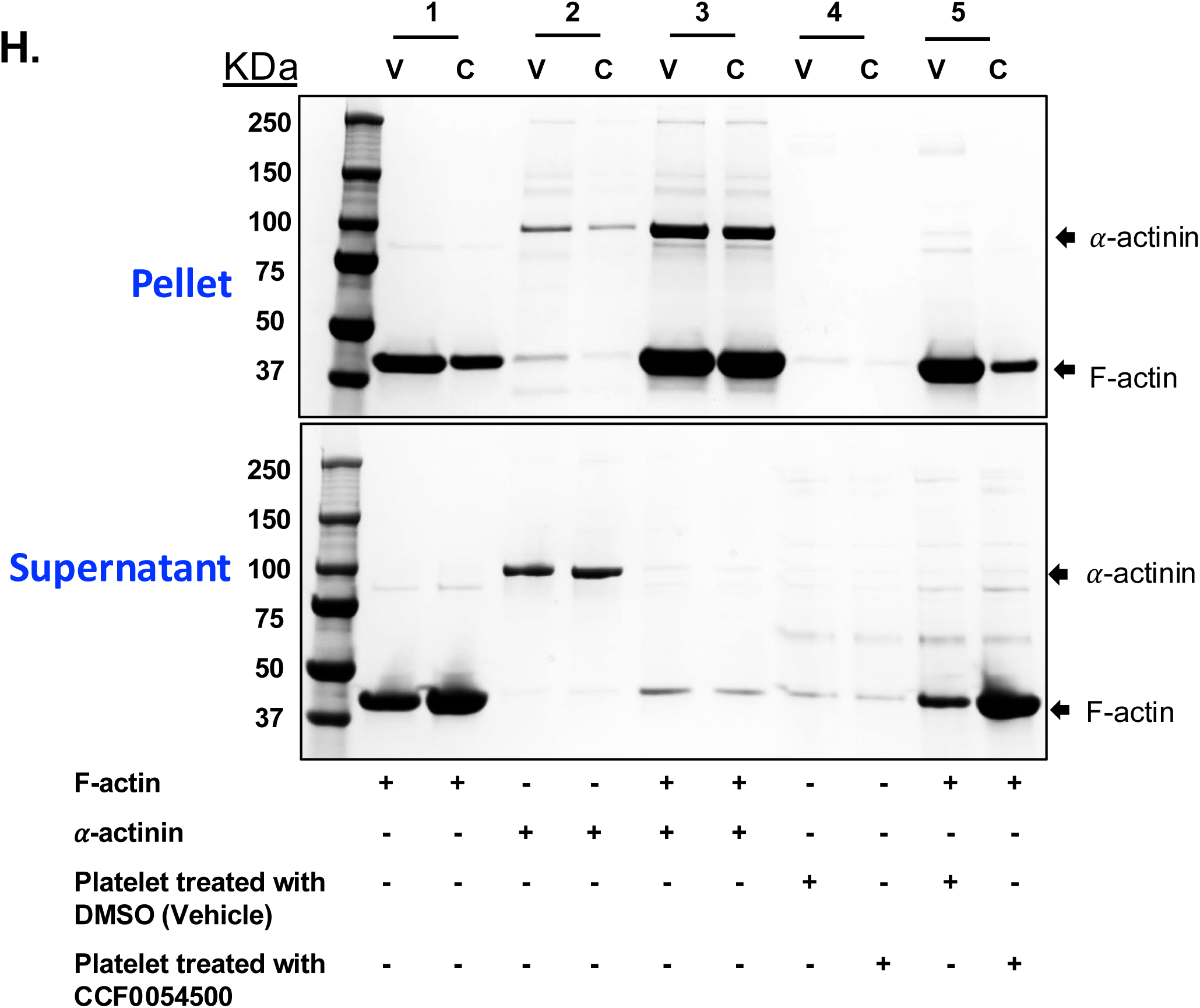
CCF0054500 rearranges the platelet actin cytoskeleton through HSP27. **(A)** Treatment of healthy human platelets with CCF0054500 vs. vehicle and unbiased phosphoproteomics and protein identification by mass spectrometry revealed drug-induced changes in the platelet actin cytoskeleton rearrangement and intracellular transport by Gene Ontology (GO) analysis with enrichment shown by Kyoto Encyclopedia of Genes and Genomes (KEGG) display. Washed platelets isolated from n=3 healthy subjects. **(B).** Treatment of human platelets with 100μM CCF0054500 and targeted phosphokinase array revealed marked phosphorylation of heat shock protein 27 (HSP27). **(C)** Densitometry of CCF0054500/vehicle-treated platelets for each phosphoprotein (n=2). **(D)** Platelets treated with CCF0054500 (0.875-112μM) or vehicle for 30 minutes and protein lysates separated by SDS-PAGE. HSP27 phosphorylation at serine 15, 78, and 82 independently validated by immunoblotting. Membranes re-probed with anti-alpha tubulin antibody to serve as a loading control. Signal intensity determined by densitometry, and values shown as the ratio of pHSP27/HSP27 for n=4-5 healthy subjects. *P* values b < 0.05 (∗) and 0.01 (∗∗). **(E)** Platelets spreading on a fibrinogen matrix illustrates actin cytoskeleton dynamics (green=filamentous actin, red=p-selectin). Surface area of healthy platelets represented as mean ± SEM, *p=0.017 by t-test for 42 random fields per subject (n=3 subjects). White scale bar=12.5µm. *P* values <0.05 (∗). **(F).** Morphological change of clot over time using washed platelets from healthy control treated with Vehicle vs CCF0054500 to illustrate cytoskeleton dynamics. Experimental tubed contained a blank (without thrombin agonist, Lane 1), thrombin (Lane 2), washed platelets treated with Vehicle (Lane 3) and CCF0054500 (Lane 4) for 30 minutes at 37°C. Image capture every 10 minute until 60 minutes. In lane 5 and 6, washed platelets were pre-treated with BVDU (HSP27 inhibitor) for 45 minutes at 37°C before vehicle or CCF0054500 for 30 minutes at 37°C, followed by CaCl_2_, Thrombin, Fibrinogen. Experimental conditions: CaCl_2_ (2mM), Thrombin (1U/mL) and Fibrinogen (1mg/mL), CCF0054500 (100µM), BVDU (60μM). **(G)** Graphical representation of clot retraction at different time points. Statistical significance evaluated by two-way ANOVA with p-value as noted. Data expressed as mean ± SEM between Vehicle vs CCF0054500 and CCF0054500 vs BVDU+CCF0054500 (n=5). *P* <0.05 (∗), <0.01 (∗∗). **(H)** Filamentous-actin (F-actin) binding assay performed on washed platelets treated with vehicle or CCF0054500. Pellet and supernatant fractions were then collected for each reaction and separated by 4-20% SDS-PAGE and stained with 0.1% Coomassie blue. Lane 1, F-actin alone. Lane 2, *α*-actinin alone. Lane 3, *α*-actinin and F-actin. Lane 4, Test sample alone. Lane 5, Test sample and F-actin. In Lane 5, in the presence of F-actin, there is less test protein in the pellet for CCF0054500-treated washed platelets than vehicle and more in supernatant, suggesting depolymerization of F-actin with CCF0054500. V= Vehicle and C=CCF0054500.

Given the established role for platelet integrin αIIbβ3 (GPIIb/IIIα) as a common mediator of platelet activation from the inside to outside and its known role as a mediator of platelet mechanotransduction and shape change ^24,25^, we evaluated platelet spreading on a fibrinogen matrix since it is known to require reorganization of the filamentous actin cytoskeleton^26^. Co-staining platelet F-actin and P-selectin revealed impaired spreading of platelets on fibrinogen following treatment with CCF0054500 compared to vehicle (**Figure 8E**). To better translate the biomechanics of platelet shape change and its impact on thrombus stability and growth, OR2L13-mediated changes in platelet shape and the kinetics of platelet F-actin reorganization were determined. Following stimulation of isolated, washed platelets by CCF0054500, we performed clot retraction studies *ex vivo*. Clot retraction is driven by the interaction between fibrin and the actin-myosin cytoskeleton of platelets, and mediated by integrin αIIbβ3 with clot retraction proportional to platelet activation and cytoskeleton rearrangement^24^. Images of washed platelets in the presence of thrombin to activate the platelet with or without CCF0054500 were taken over time (**Figure 8F**). Platelet activation causing cytoskeleton rearrangement and clot retraction was clear and progressive following thrombin stimulation (**Figure 8F**, Lane 2 vs. Lane 1), and inhibited when comparing CCF0054500 treatment to vehicle with thrombin (Lane 4 vs, Lane 3). To demonstrate the dependence of HSP27 on this process, the known HSP27 inhibitor Brivudine (BVDU)^26^ was added, showing complete abrogation of the protective effect of CCF0054500 against thrombin-mediated clot retraction (Lane 6 vs. Lane 4). The kinetics of OR2L13-agonist-mediated cytoskeleton rearrangement are summarized graphically in **Figure 8G**. Finally, to better understand the bioenergetics of F-actin depolymerization, an *in vitro* assay was used in which washed platelets from healthy subjects were treated with CCF0054500, and release of F-actin from the membranous fraction was accelerated with appearance in the soluble supernatant fraction (**Figure 8H**, lane 5 vehicle vs. control), but without any impact on globular actin (G-actin) (**Figure S8**, lane 5 vehicle vs. control) following separation of proteins by SDS-PAGE and staining with Coomassie blue. These experiments confirmed that platelet OR2L13 activation by CCF0054500 through a non-canonical pathway involving HSP27 operates by promoting F-actin depolymerization rather than preventing G-actin polymerization and this decreases platelet reactivity.

When tested for off-target effects on 78 other GPCRs, ion channels, and membrane transporters expressed in cell lines with cAMP production (Gαs-mediated), Ca^2+^ mobilization (Gαq-mediated) and nuclear hormone receptor translocation as well as limited studies that couple of ligand-gated K^+^ and Na^+^ channels (Eurofins) using CCF0054500 (0.1 nM-1000 nM) in a dose-response manner. Limited non-specific effects of CCF54500 were identified on the NAV1.5 sodium channel, the nicotinic acetyl choline receptor, the cannabinoid receptor 2, and phosphodiesterase E2D2 (**Figure S9**).

A working model is represented that may describe how CCF0054500 decreases platelet reactivity through upstream OR2L13 and downstream HSP72 phosphorylation on Ser17, Ser78, and Ser82 which promotes F-actin depolymerization of the actin cytoskeleton and prevents thrombogenic platelet granule fusion with the plasma membrane (**Figure 9**).

**Figure 9:**
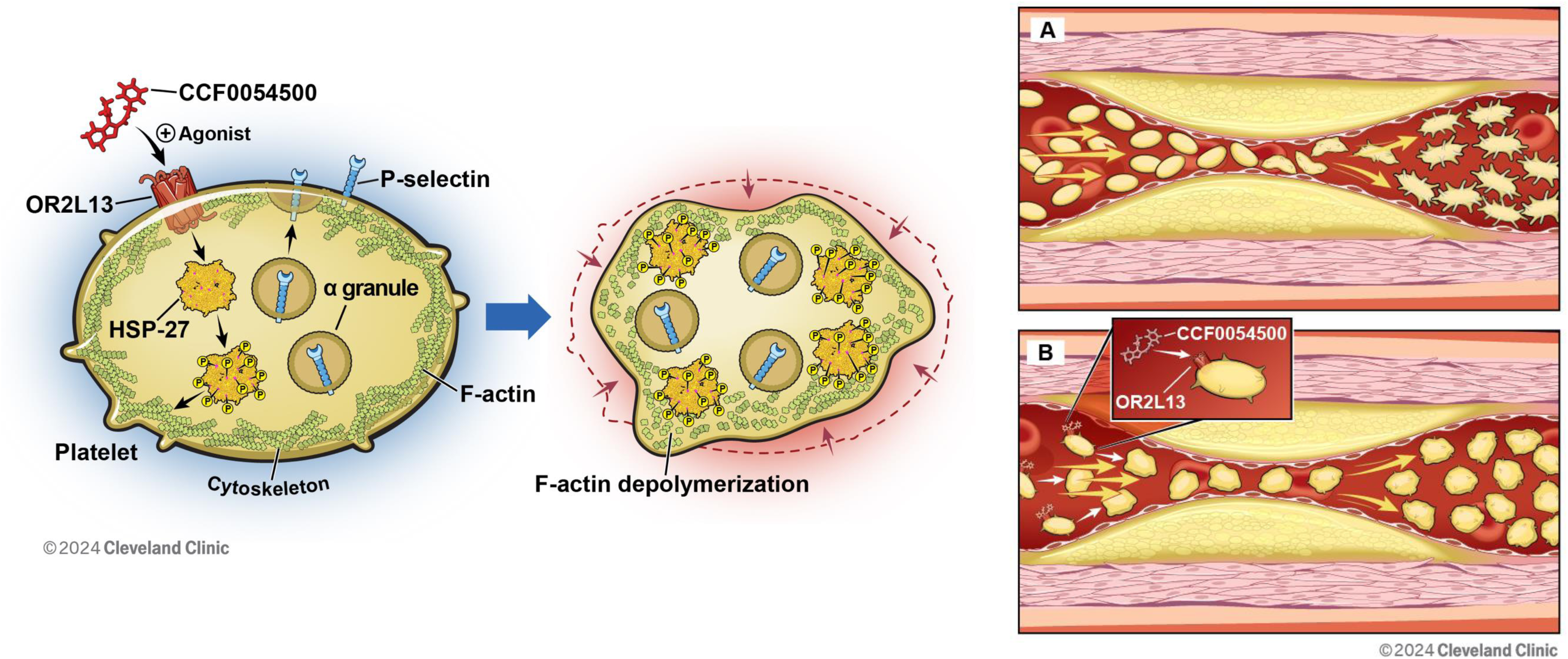
**Left:** Working model for platelet OR2L13-mediated inhibition of platelets and thrombosis through heat shock protein 27. **Right: A.** Biomechanical activation of platelets in stenosed arteries. **B.** CCF0054500 prevents biomechanical activation of platelets by altering the shape of the platelet\, rendering them less thrombogenic.

## DISCUSSION

This study identified a new class of antiplatelet drugs that operate through platelet receptor OR2L13 via a non-canonical pathway. When OR2L13 is activated by CCF0054500, platelet activation is suppressed without an impact on hemostasis or the coagulation cascade. Interestingly, CCF0054500 is equally effective at preventing thrombosis in isolated platelets, platelet rich plasma, and whole blood in both humans and mice suggesting a conserved protective cross-species pathway. This treatment strategy offers a solution to circumvent the clinical problem of HRPR and non-responsiveness to antiplatelet medications in CAD and PAD and it does so while limiting bleeding.

Dual antiplatelet therapy (DAPT) which includes aspirin and a P2Y_12_ receptor antagonist is prescribed for every patient following acute myocardial infarction according to established guidelines^4,27,28^. However, many patients—especially black patients—are resistant to the P2Y12 antagonist clopidogrel which is required for treatment of PAD or CAD^29–32^. The identification of an additional platelet GPCR target like OR2L13, coupled with a promising chemical probe like CCF0054500, offers an alternative method to circumvent a dysfunctional platelet signaling pathway in patients with clopidogrel resistance, with a poor response to aspirin therapy, or with recurrent thrombosis in spite of clopidogrel or aspirin. Prescription antiplatelet medications that include aspirin, all P2Y_12_ receptor antagonists, and especially the PAR1 antagonist vorapaxar prevent thrombosis at the expense of impairing hemostasis which is a normal physiological process to injury that limits excessive bleeding ^23,33^. Because each of these medications also impairs hemostasis and increases bleeding, this creates complications in clinical care by compelling physicians to stop antiplatelet medications that patients require to mitigate thrombosis.

In patients with both PAD and CAD, CCF0054500 attenuated all platelet surface receptor pathways that prescription drugs are targeted against, with the notable preference for its marked inhibition of GPVI. GVI is well-known to be a potent platelet signal transduction pathway for activating thrombosis without interfering with the protective mechanism of hemostasis^34–36,37^ and this may explain the mechanism of action of CCF0054500. Preliminary data from murine models *in vivo* demonstrate our OR2L13-modulating compound CCF0054500 is systemically available and inhibits thrombosis without impairing hemostasis No currently available prescription standard-of-care anti-platelet medication is capable of this.

An additional benefit of using a non-olfactory ligand for an olfactory receptor is that it overcomes several limitations of odorant olfactory ligands. For example, odorants are problematic as therapeutic agents because of their volatility and bioavailability, which limits their potential as therapeutic agents. Furthermore, odorants tend to be promiscuous, stimulating many of the approximately 400 olfactory receptors in humans leading to a lack of specificity^38,39,40^. While OR2L13 stimulation generates cAMP, a known potent inhibitor of platelets, the non-canonical pathway discovered here operates through the common downstream mediator HSP27 which appears to be the dominant mechanistic explanation for the antiplatelet effect of CCF0054500 given complete reversal of this protective effect when a HSP27-specific inhibitor is introduced. Heat shock proteins are emerging targets to abrogate thrombosis with HSP47 recently demonstrated to prevent thrombosis in immobilized humas and in hibernating bears^41^ and, indeed, HSP27 was shown previously to translocate, and associate with the actin cytoskeleton as a phosphoprotein in activated platelets which is a mechanism fully reproduced in our study ^42^. Another study demonstrated phosphorylation of HSP on Ser15, Ser78, and Ser82 in platelets impaired F-actin polymerization—a process reversed by mutation of those sites to prevent phosphorylation—though the impact on platelet activation was not assessed ^43^. Our data confirm those prior observations. Platelets adhere to extracellular matrix-coated surfaces, triggering sequential events like shape change, filopodia protrusion, and flattening. Platelets treated with CCF0054500 did not spread on a fibrinogen-coated surface, and CCF0054500 depolymerizes actin filaments. Actin polymerization in platelets allows for the formation and elongation of filopodia protrusions and lamellipodia and likely controls regulated exocytosis of thrombogenic granules which was also an observation in this study. Filopodia in platelets promote adhesion and limit blood flow, leading to platelet flattening and stable thrombus formation ^44^. Our current data shows that when HSP27 is phosphorylated at Ser15, Ser78, and Ser82, it destabilizes and depolymerizes long F-actin filaments which may limit efficient fusion of prothrombotic granules with the plasma membrane and subsequent exocytosis. This was demonstrated by the significant reduction of P-selectin release coinciding with decreased platelet aggregation. Thus, Ser15 as shown by Butt and colleagues may be a stimulus for F-actin depolymerization ^43^. Yet another study showed phosphorylated HSP27 when released from platelets promotes activation and so its presence inside platelets may prevent deactivation ^45^.

The persistence of Atheroembolism in patients with existing vascular disease remains a problem and involves persistent platelet activation^46,47^. One potentially significant finding in this study is the HRPR through GPVI is observed in patients with CAD and PAD, regardless of the antiplatelet agents prescribed. GPVI is a known mechanosensor for platelet activation^48^. CCF0054500 prevents thrombosis in blood under high shear conditions of 1500.s^−1^ and not at 600.s^−1^ in a collagen-coated microfluidics system that promotes shear-mediated platelet activation through GPVI. This is consistent with our mechanistic data that CCF0054500 depolymerizes the actin cytoskeleton and theoretically could make the platelet membrane less susceptible to external biomechanical forces that promote platelet activation. Given our previous observation that OR2L13 is trafficked in alpha granules to the platelet membrane under conditions disturbed-flow,^19^ which would typically be anticipated in patients with established CAD and PAD, this highlights an exciting additional antiplatelet property of CCF0054500 not offered by existing antiplatelet medications.

Our results indicate CCF0054500 suppresses platelet reactivity as effectively as mono-antiplatelet therapy or DAPT and it does so by interfering with downstream common pathway for platelet activation through all cell surface receptors. Patients who are prescribed clopidogrel, aspirin, or DAPT often display HRPR, which increases the risk of ischemic vascular events ^29,49^. We show in this study that HRPR in platelets from patients with CAD and PAD are effectively suppressed by CCF0054500 which appears to have few off-target effects on other GPCRs and additional common drug targets like ion channels. When given to mice, CCF0054500 has favorable exposure at low doses, prevents platelet activation and thrombosis in multiple animal models without impacting hemostasis. This makes the development of a platelet OR2L13-modulating ligand, a potential innovative alternative for antiplatelet therapy in elderly patients who have a higher incidence of HRPR with both clopidogrel and ticagrelor^49^

We acknowledge limitations of this study including an IC_50_ for CCF0054500 in the low μM range. Subtle modification of the molecular structure of CCF0054500 are underway to refine the compound and enable traditional drug-like properties. In addition, while we show parenteral delivery of CCF0054500 is an effective antithrombotic medication in mice without impacting hemostasis, oral bioavailability should be prioritized alongside thorough pharmacokinetic studies to better predict the drug behavior *in vivo*.

In conclusion, CCF0054500 is the first non-olfactory ligand identified for an olfactory receptor and inhibits thrombosis without impairing hemostasis by depolymerizing the platelet actin cytoskeleton to limits platelet shape change. Targeting platelet OR2L13 in this manner may also have the added advantage of preventing biomechanical platelet activation in pathological conditions like stenotic atherosclerotic disease or when platelets are exposed to external shear by external cardiac support devices.

## Abbreviations

(OR2L13): Olfactory Receptor 2L13
(HTS): High Throughput Screen
(cAMP): cyclic AMP
(HSP27): Heat Shock Protein 27
(HRPR): High Residual Platelet Reactivity
(GPVI): Glycoprotein VI
(TxR): Thromboxane Receptor

## Supplemental Figures

**Figure S1:**
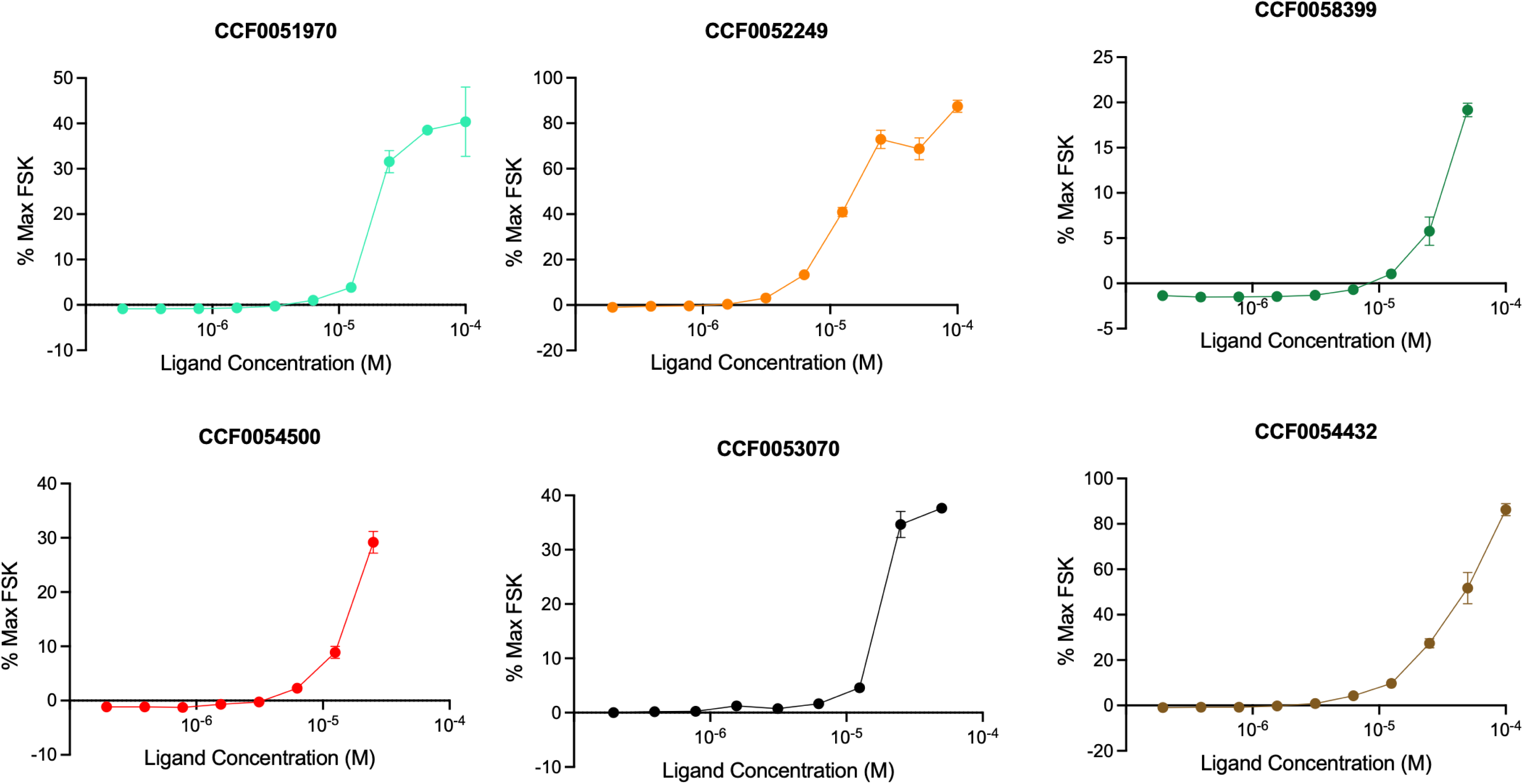
Dose response of led compounds >1.5-fold change in the 8K ligand screen in OR2L13-cAMP reporter cell line. OR2L13 activation by luciferase activity determined at different concentrations of ligand in OR2L13 HEK293 cAMP cell line. Concentrations used: 1*10^−4^ M, 5*10^−5^ M, 2.5*10^−5^ M,1.25*10^−5^ M, 6.25*10^−6^ M, 3.13*10^−6^ M, 1.568*10^−6^ M, 7.88*10^−7^ M, 3.91*10^−7^ M, 1.95*10^−7^ M.

**Figure S2:**
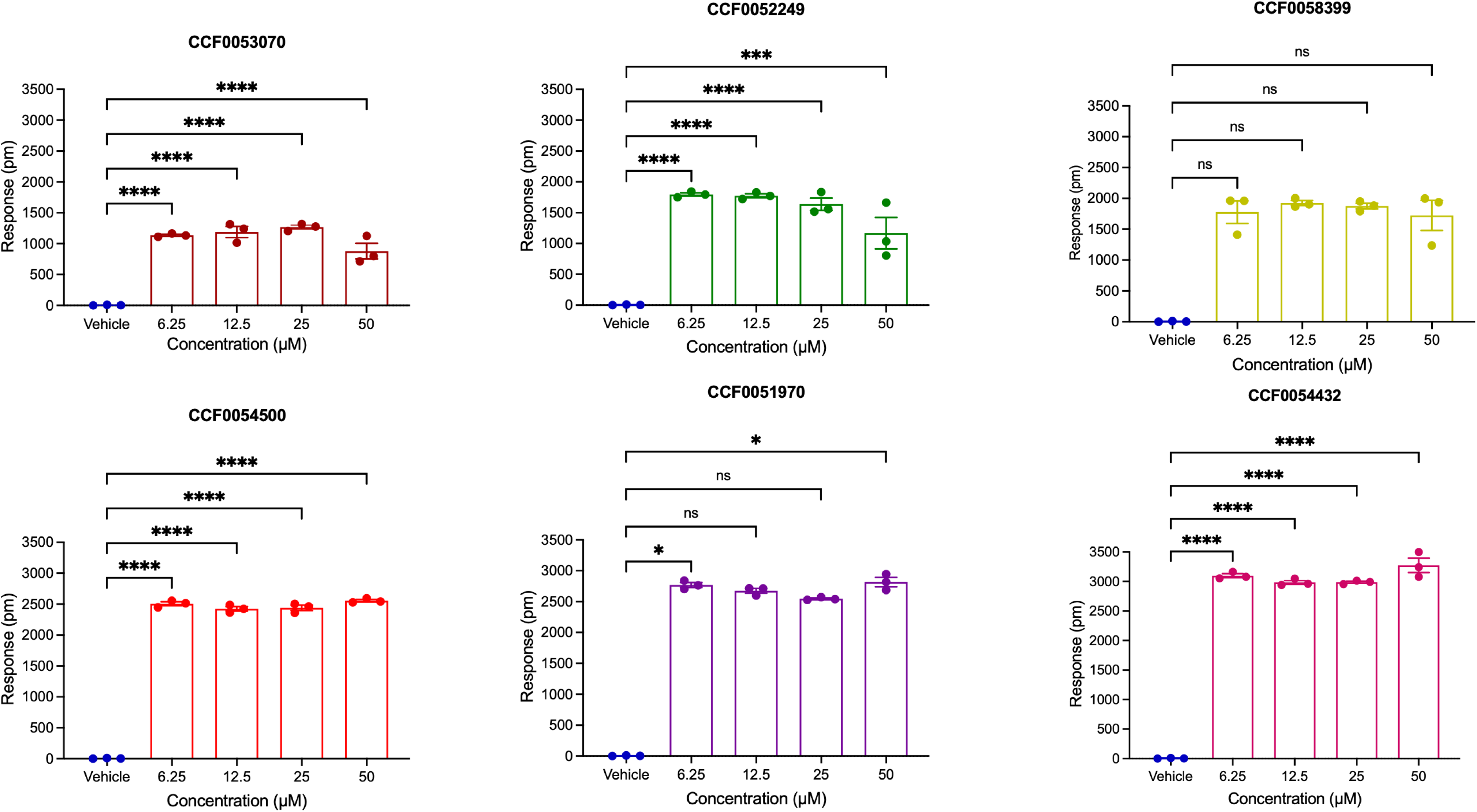
Validation of lead compounds using Dynamic Mass Redistribution: The effect of top hits at different doses (6.25μM-50μM) were validated using DMR to determine the effect of ligands on cellular responses in OR2L13 HEK293 cAMP reporter cell lines. All ligands showed displayed cellular responses in comparison to vehicle control. *P* values < 0.05 (∗), 0.01 (∗∗), 0.001 (∗∗∗), and 0.0001 (∗∗∗∗).

**Figure S3:**
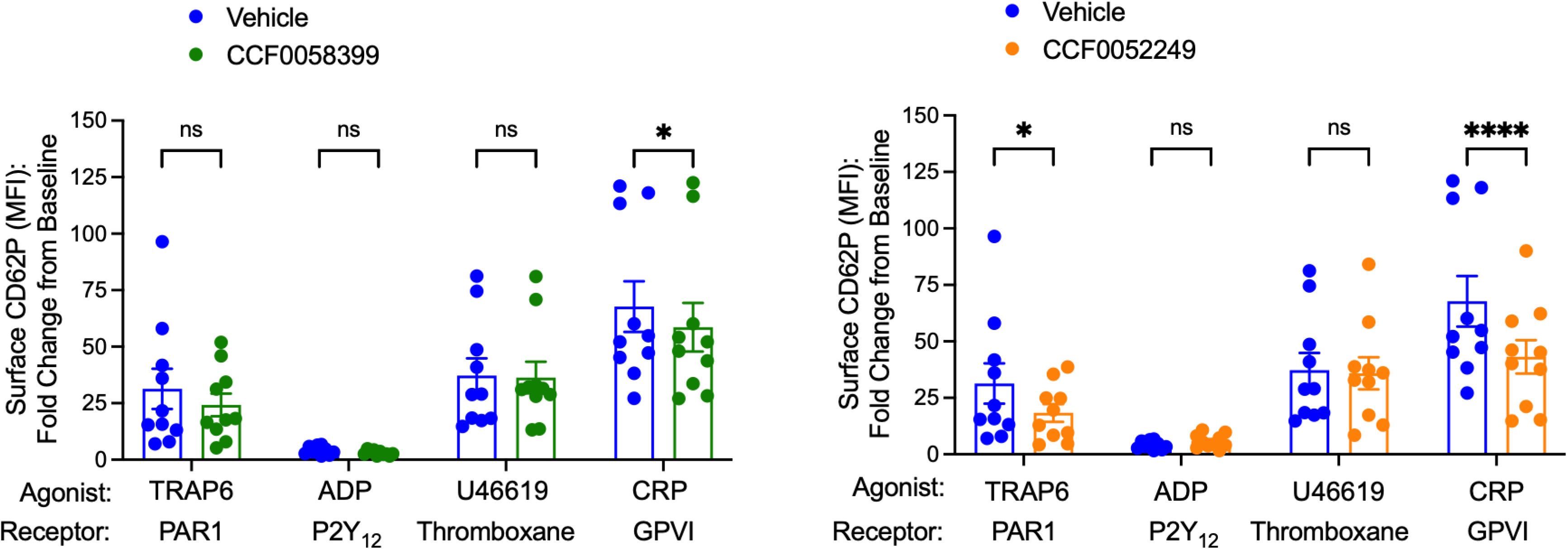
Impact of lead compounds on platelet activation. Platelet activation by alpha granule exocytosis and changes in surface P-selectin expression of washed platelets treated with agonists (CCF0058399 and CCF0052249) and vehicle for 30 minutes at 37°C. The results are expressed as Mean Fluorescence of P-selectin ± SEM (n=10, t-test). TRAP6=Thrombin Receptor Activator for Peptide 6, ADP=Adenosine Diphosphate, CRP=Collagen-Related Peptide. *P* <0.05 (∗), <0.01 (∗∗), <0.001 (∗∗∗), and <0.0001 (∗∗∗∗).

**Figure S4:**
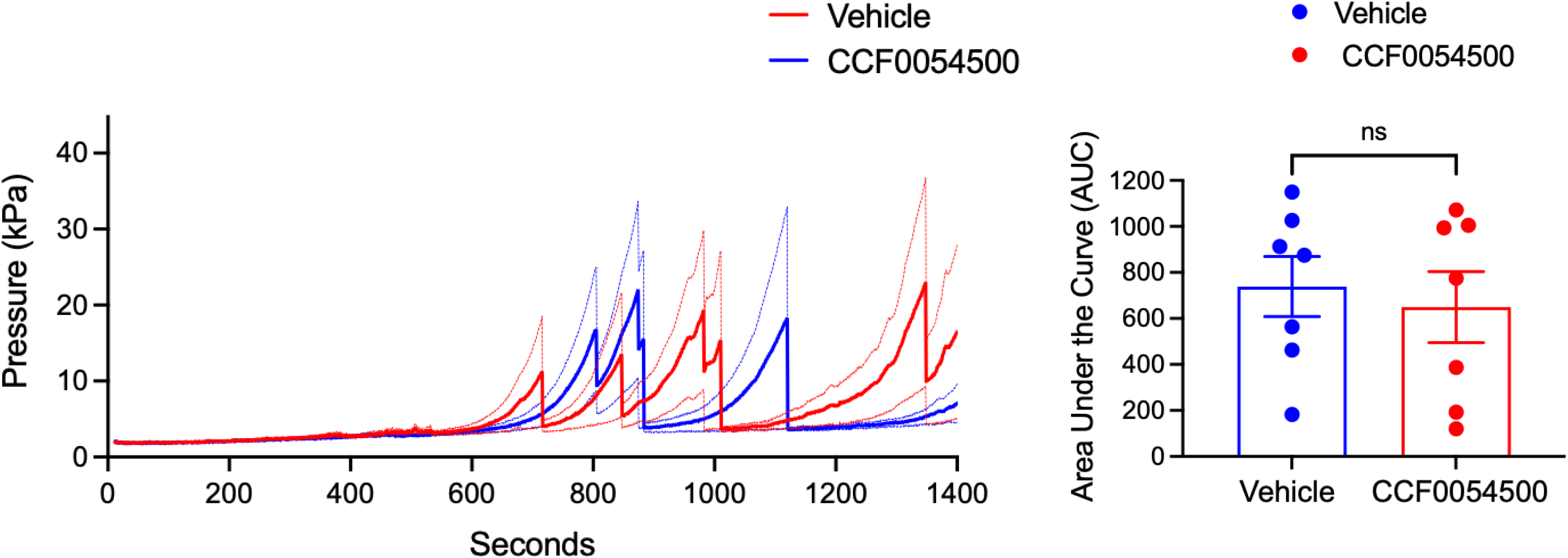
Impact of non-olfactory OR2L13 agonist lead compound on biomechanical platelet activation under low arterial stress. (A) T-TAS® (Total Thrombus-formation Analysis System) evaluates thrombosis of whole blood under lower arterial shear stress conditions (600 Sec^−1^) as a function of time and peak pressure. CCF0054500 does not change occlusion by microfluidics under low shear stress (n=7 healthy volunteers) also represented with. (B) Summary data represented as area Under the Curve. Data are represented as mean ± SEM, t-test.

**Figure S5:**
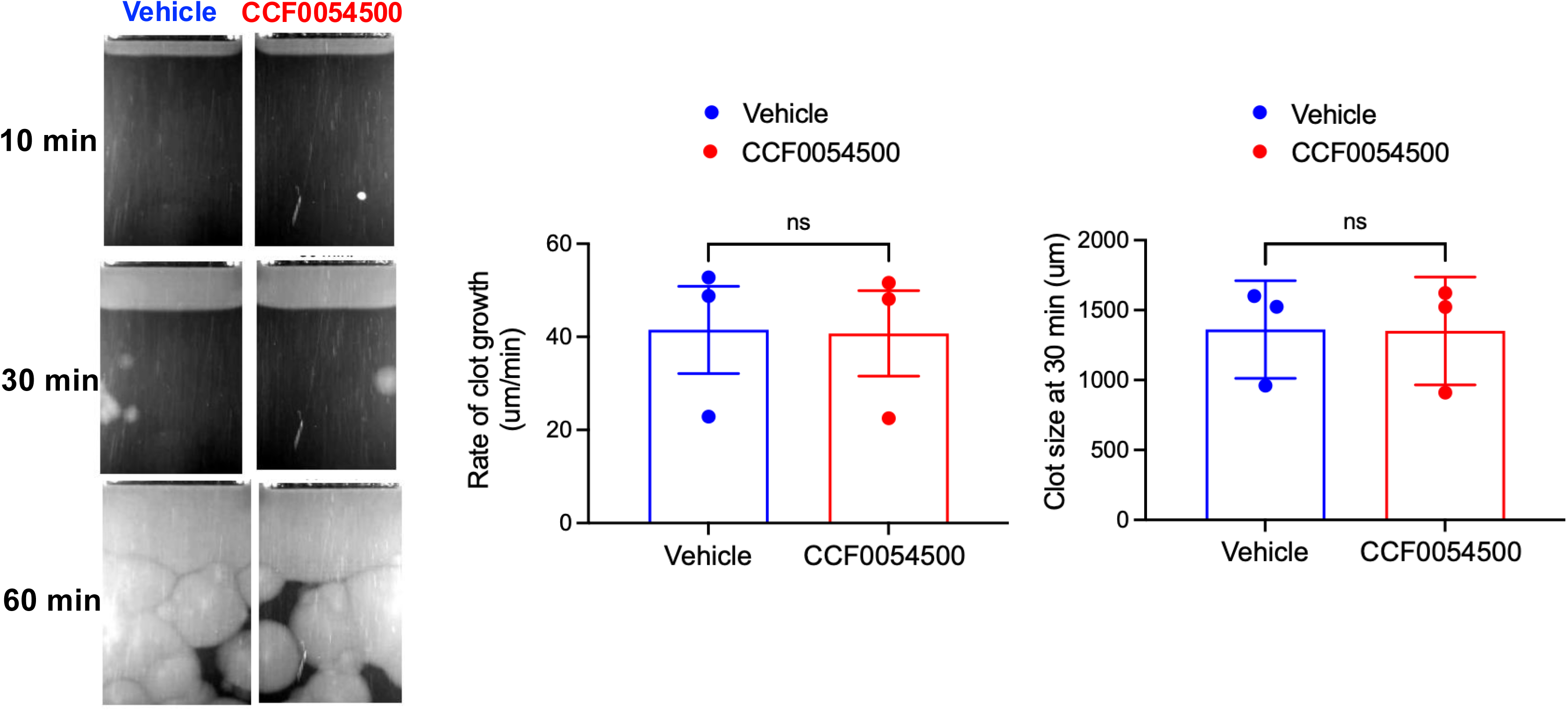
No effect of CCF0054500 on the coagulation cascade. CCF0054500 does not impact the coagulation cascade tested through Thrombodynamics ® analysis of platelet-deplete plasma. There is no difference in clot size or rate of growth of fibrin formation when treated with CCF0054500 compared with Vehicle (n=3).

**Figure S6:**
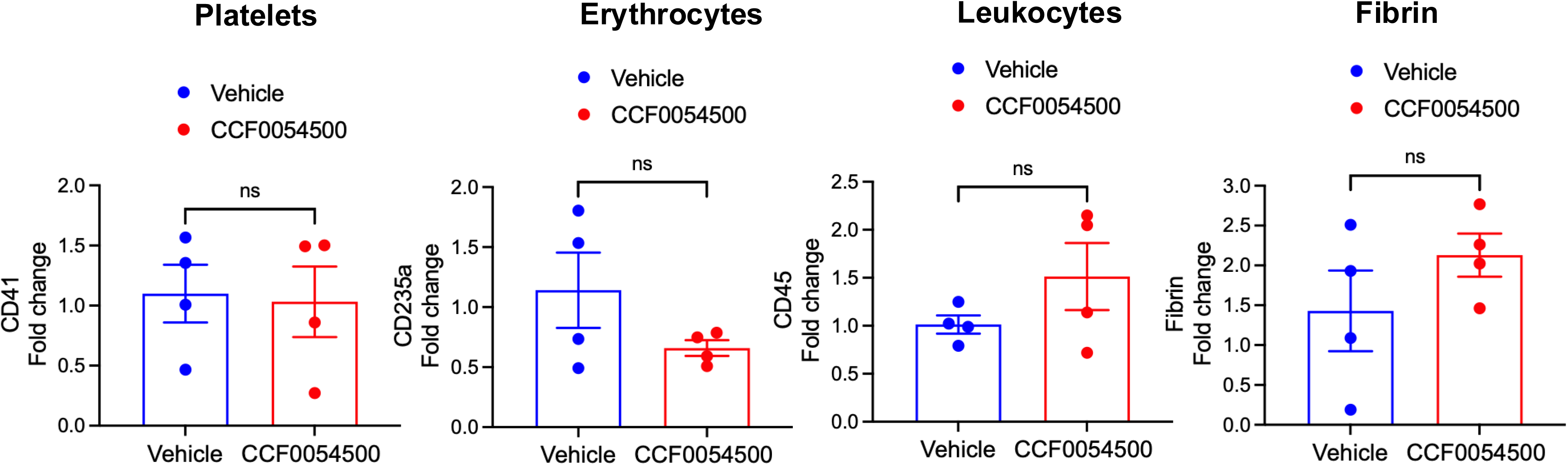
CCF0054500 does not change the composition of the cells in thrombus by IVC stasis: RNA isolated from thrombus treated with CCF0054500 or vehicle. Cell populations assessed by cell-specific markers: Platelets (CD41), Erythrocytes (CD235a), Leukocytes (CD45) and fibrin were tested using qPCR in the thrombus and there was no difference in these two groups. Data is presented as mean ± SEM, n=4 thrombi (t-test).

**Figure S7:**
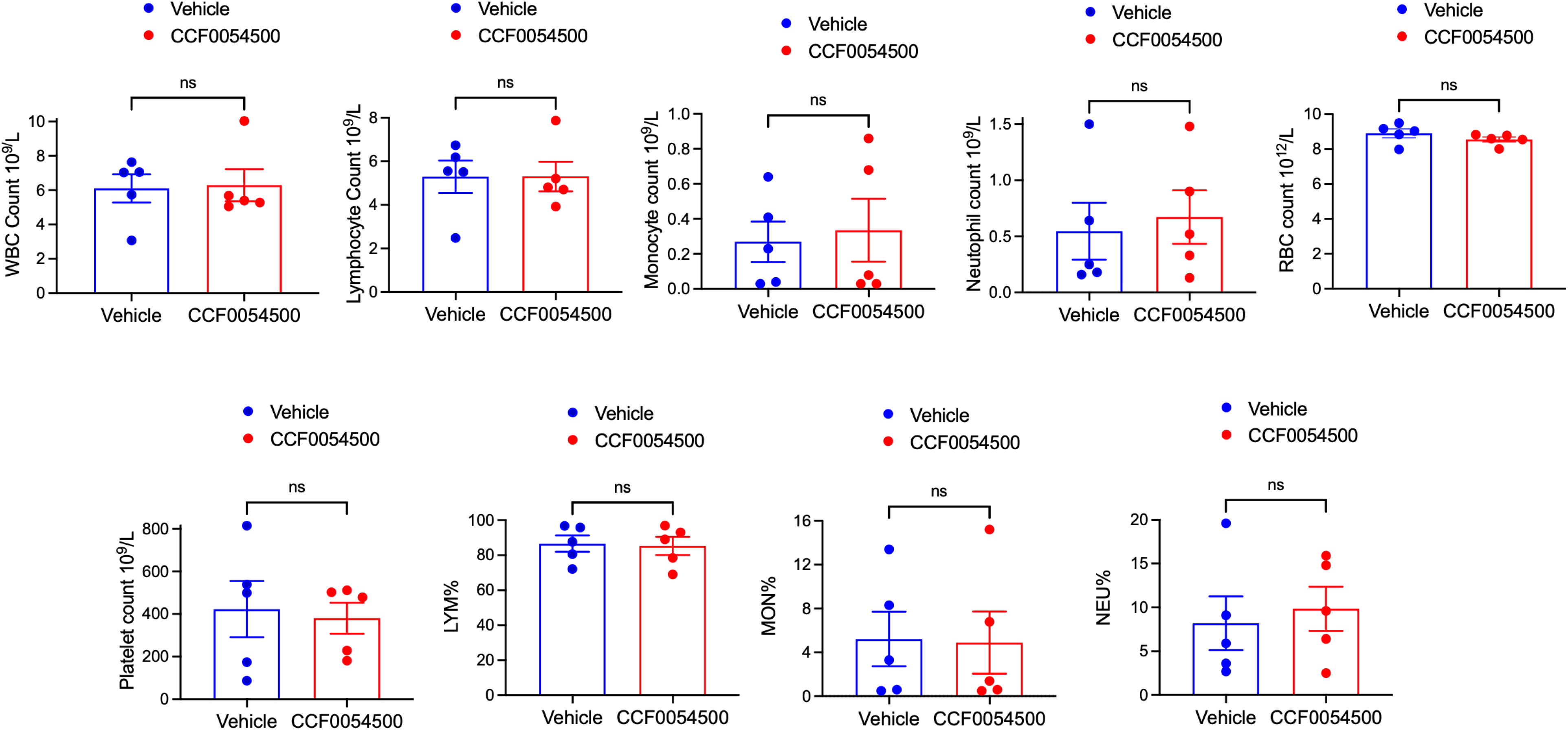
CCF0054500 does not change circulating cells in the blood of mice. No effect of CCF0054500 (5mg/kg/day) on thrombopoiesis, leukopoiesis, or hematopoiesis. RBC=red blood cells. WBC=white blood cells.

**Figure S8:**
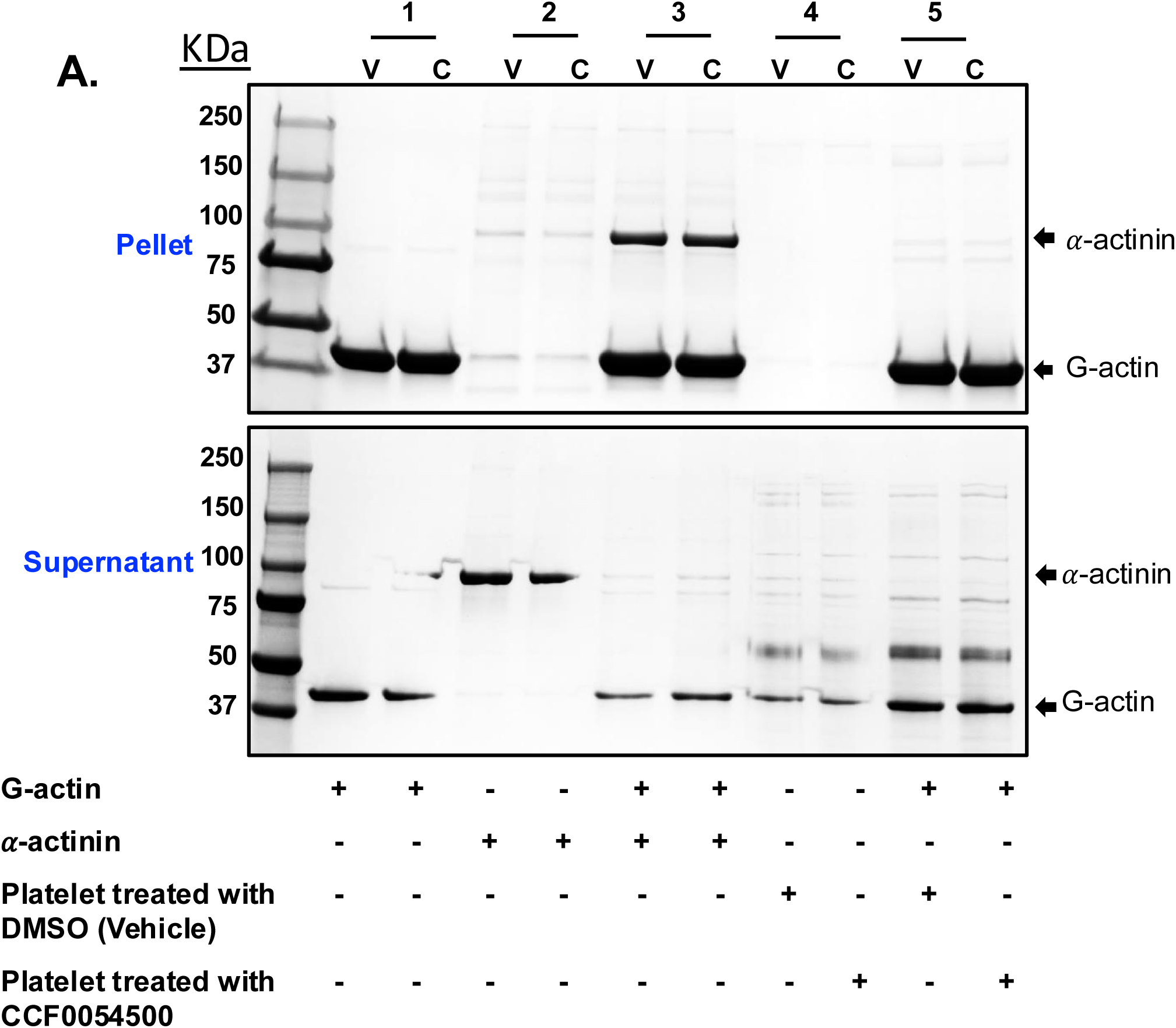
CCF0054500 does not impact the platelet cytoskeleton through HSP27 by G-actin. (A) An G-actin binding assay was performed on washed platelets treated with vehicle or 100µM CCF0054500 according to method 1 of Actin Binding Protein Biochem kit from Cytoskeleton, Inc. Samples of the Pellet and supernatant fractions were collected for each reaction and were separated on 4-20% SDS-gel and stained with 0.1% Coomassie blue. Lane 1, G-actin alone. Lane 2, *α*-actinin alone. Lane 3, *α*-actinin and F-actin. Lane 4, Test sample alone. Lane 5, Test sample and F-actin. V= Vehicle and C=CCF0054500. In Lane 5, in the presence of G-actin, there is less test protein in the pellet for CCF0054500 treated washed platelets than vehicle and more in supernatant, suggesting depolymerization of G-actin with CCF0054500.

**Figure S9:**
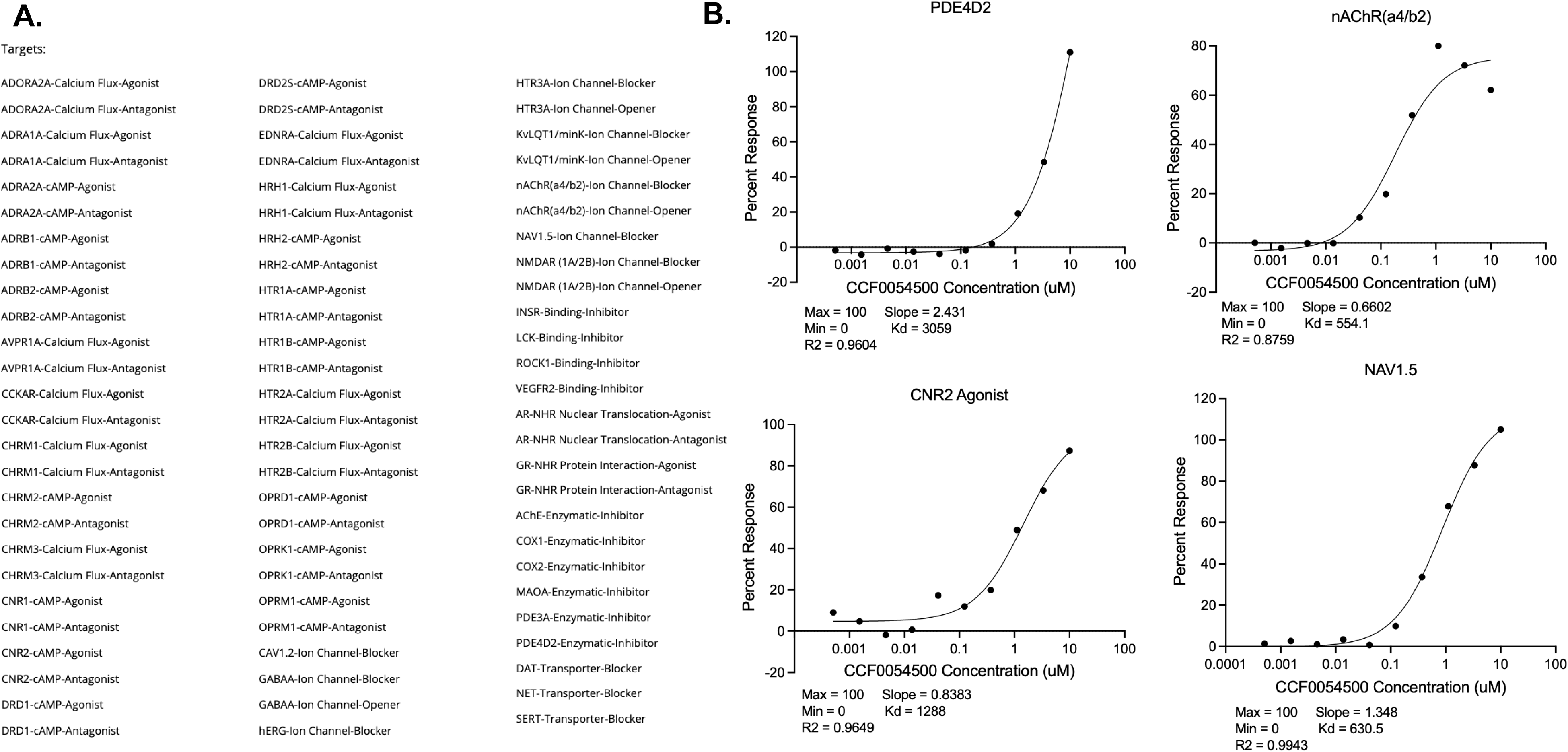
Off-target effects of CCF0054500. (A) Eurofins off-target analysis of CCF0054500 using cell lines expressing GPCRs, nuclear receptors, ion channels and enzymes. 78 Assays tested are indicated above. Off-target effects were noted for a sodium channel, a phosphodiesterase isoform, and the nicotinic acetyl choline receptor, and cannabinoid Receptor 2 (log dose-response curves indicated). All other targets were negative for cross-reactivity. (B). Accompanying Excel file is uploaded with raw pharmacological data.

## ACKNOWLEDGEMENTS

This work was supported by the National Institutes of Health grant 26512550 (SJC), NCAI 1U54HL119810-07 (SS, SJC, JVP), a SPARK Catalyst Award from Cleveland Clinic (SJC), K99HL164888 (M.Y.), and the American Society of Hematology Scholar Award (M.Y.). Screening, hit confirmation, and compound re-synthesis was provided by the Center for Therapeutics Discovery (C3TD), Cleveland Clinic, Cleveland, OH 44195. We thank C3TD staff for insightful discussions and assistance with assay optimization and execution.

